# Acclimation of *Chlamydomonas reinhardtii* to Nitric Oxide Stress Related to Respiratory Burst Oxidase-Like 2

**DOI:** 10.1101/2021.03.30.437739

**Authors:** Eva YuHua Kuo, Tse-Min Lee

## Abstract

The acclimation mechanism of *Chlamydomonas reinhardtii* to nitric oxide (NO) was studied by exposure to *S*-nitroso-*N*-acetylpenicillamine (SNAP), a NO donor. Treatment with 0.1 or 0.3 mM SNAP transiently inhibited photosynthesis within 1 h, followed by a recovery without growth impairment, while 1.0 mM SNAP treatment caused irreversible photosynthesis inhibition and mortality. The SNAP effects are avoided in the presence of the NO scavenger, 2-(4-carboxyphenyl)-4,4,5,5-tetramethylimidazoline-l-oxyl-3-oxide (cPTIO). RNA-seq, qPCR, and biochemical analyses were conducted to decode the metabolic shifts under sub-lethal NO stress by exposure to 0.3 mM SNAP in the presence or absence of 0.4 mM cPTIO. These findings revealed that the acclimation to NO stress comprises a temporally orchestrated implementation of metabolic processes: 1. trigger of NO scavenging elements to reduce NO level; 2. prevention of photo-oxidative risk through photosynthesis inhibition and antioxidant defense system induction; 3. acclimation to nitrogen and sulfur shortage; 4. degradation of damaged proteins through protein trafficking machinery (ubiquitin, SNARE, and autophagy) and molecular chaperone system for dynamic regulation of protein homeostasis. NO increased NADPH oxidase activity and respiratory burst oxidase-like 2 (RBOL2) transcript abundance, which were not observed in the *rbol2* insertion mutant. Changes in gene expression in the *rbol2* mutant and increased mortality under NO stress demonstrate that NADPH oxidase (RBOL2) is involved in the modulation of some acclimation processes (NO scavenging, antioxidant defense system, autophagy, and heat shock proteins) for *Chlamydomonas* to cope with NO stress. Our findings provide insight into the molecular events underlying acclimation mechanisms in *Chlamydomonas* to sub-lethal NO stress.

**One-sentence Summary:** Acclimation machinery is triggered in *Chlamydomonas reinhardtii* cells against sub-lethal nitric oxide stress.

## INTRODUCTION

Nitric oxide (NO) is a crucial signaling molecule in diverse physiological processes in plants (Durner et al., 1999), including in the green alga *Chlamydomonas*, in which NO regulates many physiological processes and stress responses such as the remodeling of chloroplast proteins by the degradation of thylakoid cytochrome *b*_6_*f* complex and stroma ribulose-1,5-bisphosphate carboxylase/oxygenase (Rubisco) via FtsH and Clp chloroplast proteases under nitrogen (Wei et al., 2014) or sulfur starvation (de Mia et al., 2019). In *Chlamydomonas*, NO is also involved in cell death induced by ethylene and mastoparan (Yordanova et al., 2010), induction of oxidative stress under extreme high light (VHL, 3,000 μmol·m^−2^·s^−1^) (Chang et al., 2013), interaction of NO with hydrogen peroxide (H_2_O_2_) for high light stress-induced autophagy and cell death (Kuo et al., 2020a), proline biosynthesis under copper stress (Zhang et al., 2008), and responses to salt stress (Chen et al., 2016). NO is responsible for the regulation of nitrogen assimilation by repressing the expression of nitrate reductase as well as nitrate and ammonium transporters (de Montaigu et al., 2010) and their enzyme activities (Sanz-Luque et al., 2013; Calatrava et al., 2017). Mitochondrial respiration by upregulation of alternative oxidase 1 is modulated by NO in *Chlamydomonas* (Zalutskaya et al., 2017).

Information is still lacking on the comprehensive overview of the transcriptional regulation of *C*. *reinhardtii* in response to NO stress. Therefore, in the present study, *C. reinhardtii* cells were treated with different concentrations (0.1, 0.3, and 1.0 mM) of the NO donor *S*-nitroso-*N*-acetylpenicillamine (SNAP) in the presence or absence of an NO scavenger, 2-(4-carboxyphenyl)-4,4,5,5-tetramethylimidazoline-l-oxyl-3-oxide (cPTIO) (Mur et al., 2011). This study provides new insight into the acclimation mechanisms against NO stress in *Chlamydomonas*.

## RESULTS

### Physiological and Transcriptomic Changes in Response to NO Burst

The mechanisms that allow for the acclimation of *C*. *reinhardtrii* cells to short-term NO burst were examined. The SNAP concentration and treatment duration were carefully chosen to allow for the monitoring of the short-term response to NO rather than cell death. Using a cell permeable NO-sensitive fluorescent dye, the fluorescence emitted from the cells (Fig. 1A) rapidly increased 0.5 h after SNAP treatment and reached a plateau after 1 h (Fig. 1B); the increase in fluorescence can be inhibited in the presence of 0.4 mM cPTIO. Both the 0.1 and 0.3 mM SNAP treatments did not affect cell viability (Fig. 1C) and growth (Fig. 1D), whereas the 1.0 mM SNAP treatment resulted in cell mortality and bleaching (Fig. 1E). The concentrations of chlorophyll *a* (Supplemental Fig. S1A), chlorophyll *b* (Supplemental Fig. S1B), and carotenoids (Supplemental Fig. S1C) were not influenced by either 0.1 or 0.3 mM SNAP treatments; however, a decrease in chlorophyll *a*, chlorophyll *b*, and carotenoids 6 h after treatment with 1.0 mM SNAP was observed. Photosynthesis is relatively sensitive to NO burst, as reflected by a transient decrease in *F*_v_′/*F*_m_′ (Fig. 2A), *F*_v_/*F*_m_ (Fig. 2B), and the O_2_ evolution rate (Fig. 2C) 0.5 h after SNAP treatment; these photosynthesis-related factors then recovered after 3–5 h in both the 0.1 and 0.3 mM SNAP treatments but not in the 1.0 mM SNAP treatment. This inhibition of photosynthesis-related factors can be suppressed in the presence of 0.4 mM cPTIO. Based on fluorescence induction kinetics (OJIP curve), the cells treated with 0.1 or 0.3 mM SNAP transiently lost both the PSII acceptor-side (*F*_J_-*F*_o_ and *F*_I_-*F*_J_) and donor-side (*F*_P_-*F*_I_ and *F*_v_/*F*_o_) electron transfer ability, whereas 1.0 mM SNAP treatment caused irreversible inhibition, which was prevented in the presence of 0.4 mM cPTIO (Supplemental Table S1). The SNAP treatments decreased the production of superoxide anion radical (O_2_^.−^) and H_2_O_2_; the production of singlet oxygen (^1^O_2_) was significantly increased by 1.0 mM SNAP treatment (Supplemental Fig. S1D-F). Treatment with 1.0 mM SNAP also caused significant lipid peroxidation (Supplemental Fig. S1G). Overall, these results demonstrate two stages of response in *C*. *reinhardtii* cells in the sub-lethal NO challenge: I. NO stress together with the significant NO burst caused a sharp decline in photosynthetic activity 1 h after SNAP treatment and II. the recovery period. Therefore, *Chlamydomonas* cells treated with 0.3 mM SNAP for 1 h in the presence or absence of 0.4 mM cPTIO were used for transcriptomic analyses.

**Figure 1.**
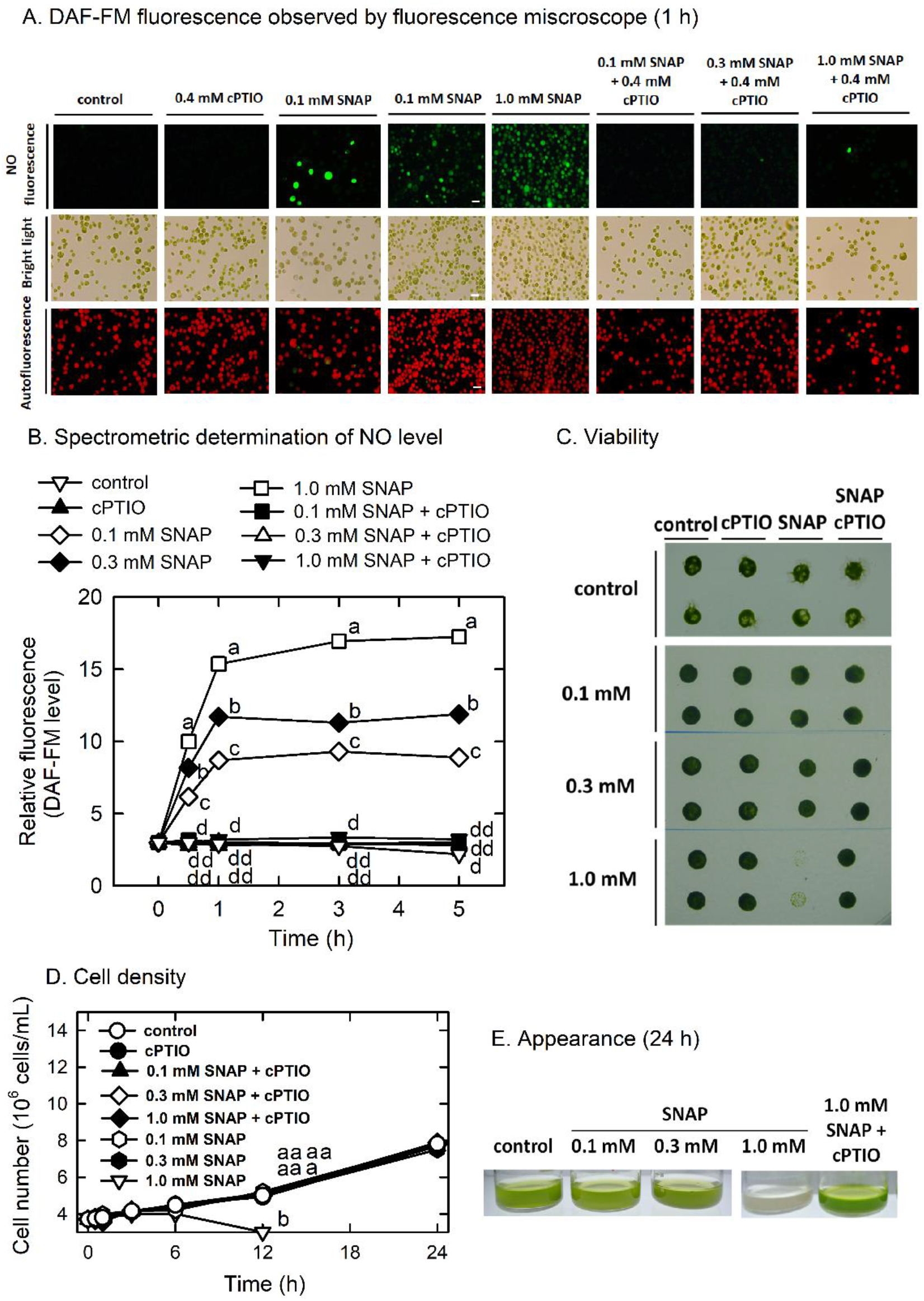
Microscopic observation (A) and spectrophotometric determination (B) of nitric oxide fluorescence (DAF-FM), cell viability (C), cell growth (D), and appearance (E) in *Chlamydomonas reinhardtii* in response to 0.1, 0.3, or 1.0 mM SNAP in the presence or absence of 0.4 mM cPTIO. Data are expressed as the mean ± SD (n = 3). Different symbols indicate significant differences between treatments (Scheffe’s test, *P* < 0.05).

**Figure 2.**
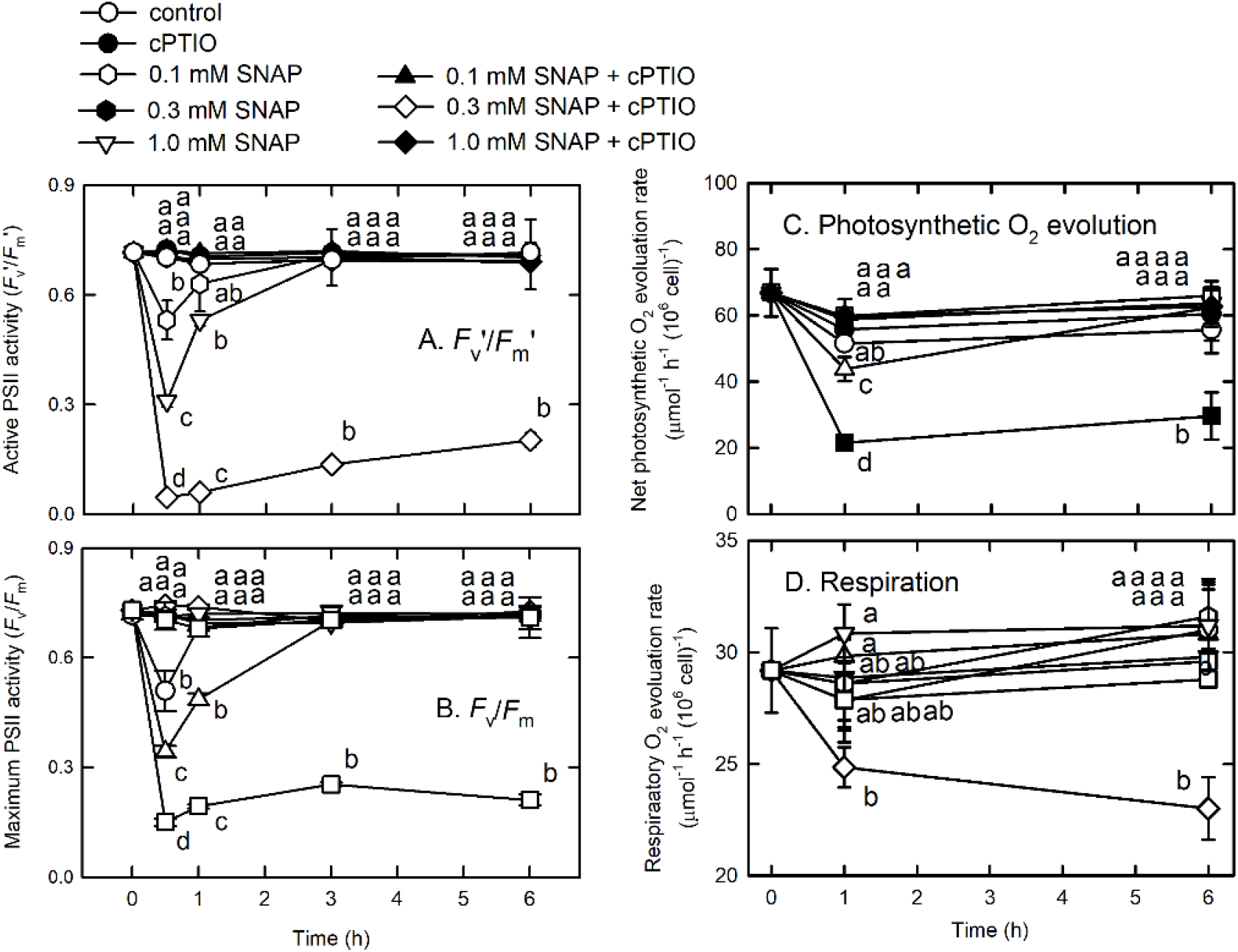
The active (*F*_v_′/*F*_m_′) (A) and maximum (*F*_v_/*F*_m_) (B) PSII activity, photosynthetic O2 evolution rate (C), and respiration rate (D) in *C. reinhardtii* after exposure to 0.1, 0.3, or 1.0 mM SNAP in the presence or absence of 0.4 mM cPTIO. Data are expressed as the mean ± SD (n = 3). Different symbols indicate significant differences between treatments (Scheffe’s test, *P* < 0.05).

The results of the transcriptomic analysis (Supplemental Tables S2 and S3) showed that 1,012 significant DEGs were regulated by NO (1.2 log_2_FC, *P*-value of log_2_FC ≤ 0.05) (Supplemental Table S4). Following analysis using the MapMan display mode, cellular response overview (Supplemental Fig. S2), proteasome and autophagy (Supplemental Fig. S3), photosynthesis (electron transport, Calvin cycle, and photorespiration) (Supplemental Fig. S4), and the tetrapyrrole pathway (Supplemental Fig. S5) were affected under NO stress. Using the Blast2GO suite, the analysis of all the functional DEGs and unigenes identified 45 GO terms (score ≤ 0.05) that can be assigned to 463 upregulated genes (45.75%), with 13 biological process, 30 molecular function, and two cellular component GO terms (Supplemental Fig. S6A and Table S5A). One hundred fifty-seven GO terms for 549 downregulated genes with 337 known function genes (54.25%) were classified into 35 GO terms belonging to biological process, 112 to molecular function, and eight to cellular component (Supplemental Fig. S6B and Table S5B). The genes associated with amino acid catabolism, the ubiquitin-proteasome system, and the antioxidant defense system are upregulated, while those related to transcriptional and translational regulation are downregulated by NO burst (Supplemental Tables S4 and S5).

We also found that the genes associated with photosynthesis are downregulated by NO burst (Supplemental Tables S7). NO decreased the transcript abundances of lumenal PsbP-like protein (PSBP4, Cre03.g182551.t1.2), which is linked to the stability of photosystem II complex assembly (Ido et al., 2014); Lhl2 (Cre08.g384650.t1.2) belonging to high light-induced protein (Teramoto et al., 2004), Lhl3 (Cre03.g199535.t1.1); pre-apoplastocyanin (PCY1, Cre03.g182551.t1.2), cytochrome c_6A_ (CYC4, Cre16.g670950.t1.2) acting as electron transfer between cytochrome *b*_6_*f* complex and photosystem I (Marcaida et al., 2006); CHLH (Cre07.g325500.t1.1); CHLI1 (Cre06.g306300.t1.2); CHLI2 (Cre12.g510800.t1.2), CHLD (Cre05.g242000.t1.2); and GENOMES UNCOUPLED 4 (GUN4, Cre05.g246800.t1.2). In contrast, NO increased the transcript abundance of chlorophyll *a*/*b* binding protein (LHCBM9, Cre06.g284200.t1.2), Lhl1 (Cre08.g384650.t1.2), and Lhl4 (Cre17.g740950.t1.2) (Fig. 3). For the Calvin cycle, NO also decreased the transcript abundance of the Rubisco small subunit 1 (RBCS1, Cre02.g120100.t1.2), ribulose phosphate-3-epimerase (RPE2, Cre02.g116450.t1.2), phosphoglycerate kinase (PGK1, Cre11.g467770.t1.1), glyceraldehyde-3-phosphate dehydrogenase (GAP4, Cre12.g556600.t1.2), triose phosphate isomerase (TPI1, Cre01.g029300.t1.2), sugar bisphosphatase (SBP2, Cre17.g699600.t1.2), a pentatrichopeptide repeat protein that stabilizes Rubisco large subunit mRNA (PPR2, Cre06.g298300.t1.1), a small protein for phosphoribulokinase deactivation (Howard et al., 2008) (CP12, Cre08.g380250.t1.2), Rubisco large (RMT1, Cre16.g661350. t1.2 and a similar one, Cre16.g649700.t1.1) and small (RMT2, Cre12.g524500.t1.2) subunit N-methyltransferase, and phosphoglycolate phosphatase (PGP3, Cre06.g271400.t1.1) in the photorespiration (Fig. 4). In contrast, the transcript abundance of Rubisco activase-like protein (RCA2, Cre06.g298300.t1.1) was increased by NO (Fig. 4H). The NO-induced changes in gene expression occurred 1 h after NO exposure, recovered to the same levels as the control after 3 h, and the changes at 1 h can be inhibited by the presence of cPTIO.

**Figure 3.**
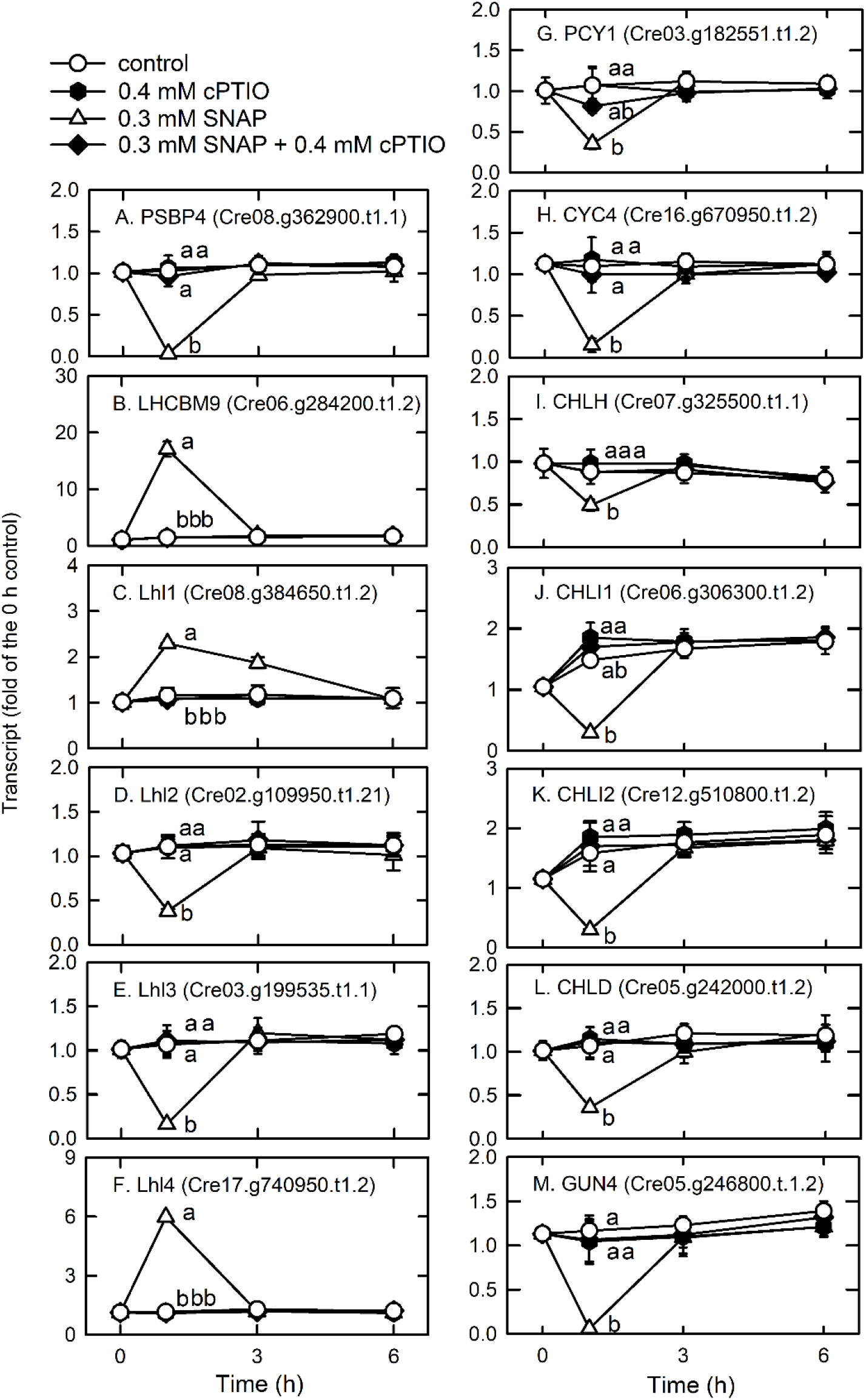
Time-course changes in the transcript abundances of PSBP4 (A), LHCBM9 (B), Lhl1 (C), Lhl2 (D), Lhl3 (E), Lhl4 (F), PCY1 (G), CYC4 (H), CHLH (I), CHLI1 (J), CHLI2 (K), CHLD (L), and GUN4 (M) in *C. reinhardtii* upon exposure to 0.3 mM SNAP in the presence or absence of 0.4 mM cPTIO. Data are expressed as the mean ± SD (n = 3). Different symbols indicate significant differences between treatments (Scheffe’s test, *P* < 0.05).

**Figure 4.**
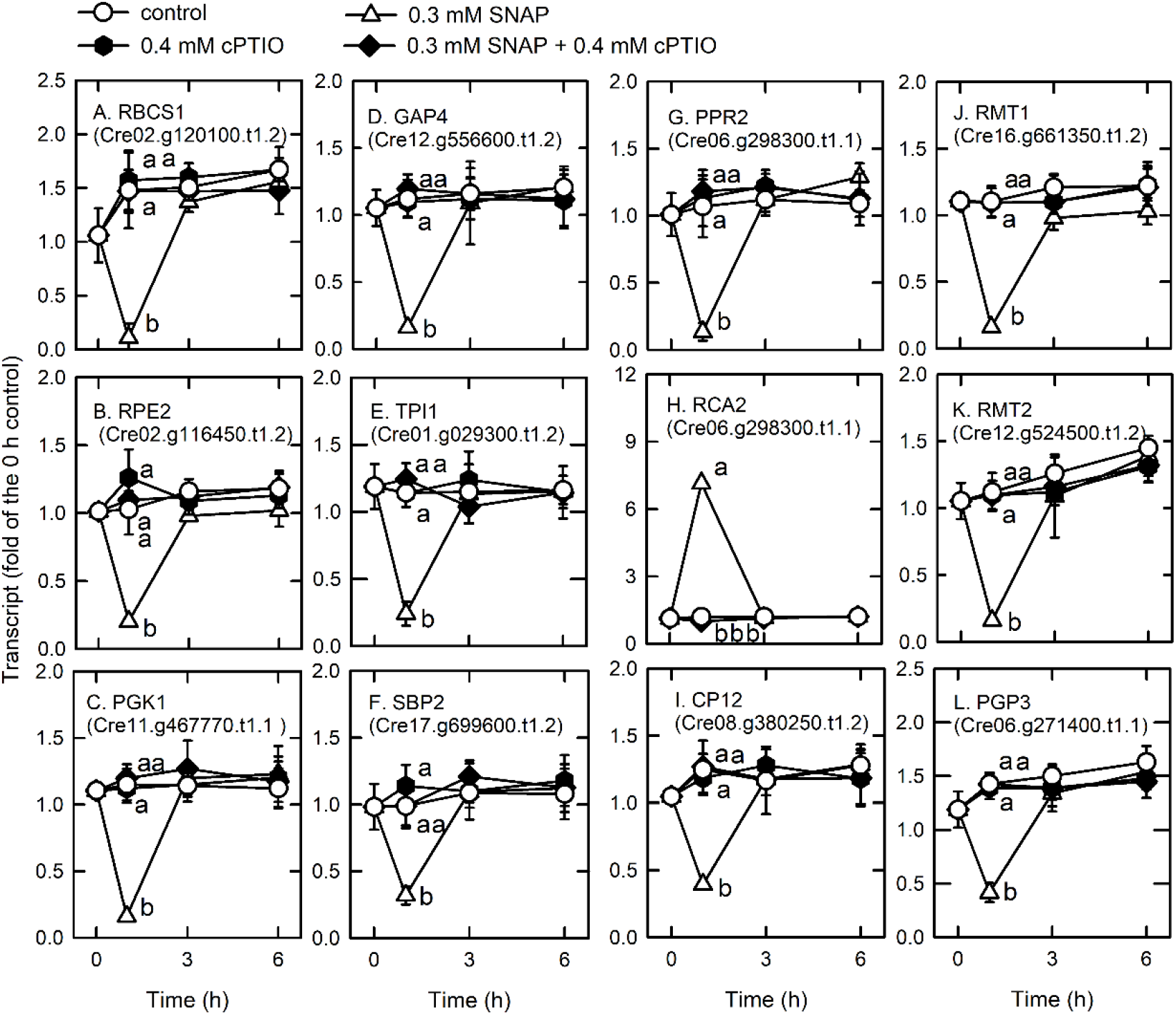
Time-course changes in the transcript abundances of RBCS1 (A), RPE2 (B), PGK1 (C), GAP4 (D), TPI1 (E), SBP2 (F), PPR2 (G), RCA2 (H), CP12 (I), RMT1 (J), RMT2 (K), and PGP3 (L) in *C. reinhardtii* after exposure to 0.3 mM SNAP in the presence or absence of 0.4 mM cPTIO. Data are expressed as the mean ± SD (n = 3). Different symbols indicate significant differences between treatments (Scheffe’s test, *P* < 0.05).

### NO Decreases Nitrogen and Sulfur Availability

The transcript abundances of the ammonium transporters AMT3 (Cre06.g293051.t1.1) (Fig. 5A), AMT6 (Cre07.g355650.t1.1) (Fig. 5B), and AMT7 (Cre02.g111050.t1.1) (Fig. 5C), and the ammonium assimilation enzymes, glutamate synthase (GSN1, Cre13.g592200.t1.2) (Fig. 5D) and glutamate synthetase (GLN1, Cre02.g113200.t1.1) (Fig. 5E) decreased 1 h after NO treatment and then recovered, while that of glutamate dehydrogenase (GDH1, Cre09.g388800.t1.2) increased after 1 h (Fig. 5F).

**Figure 5.**
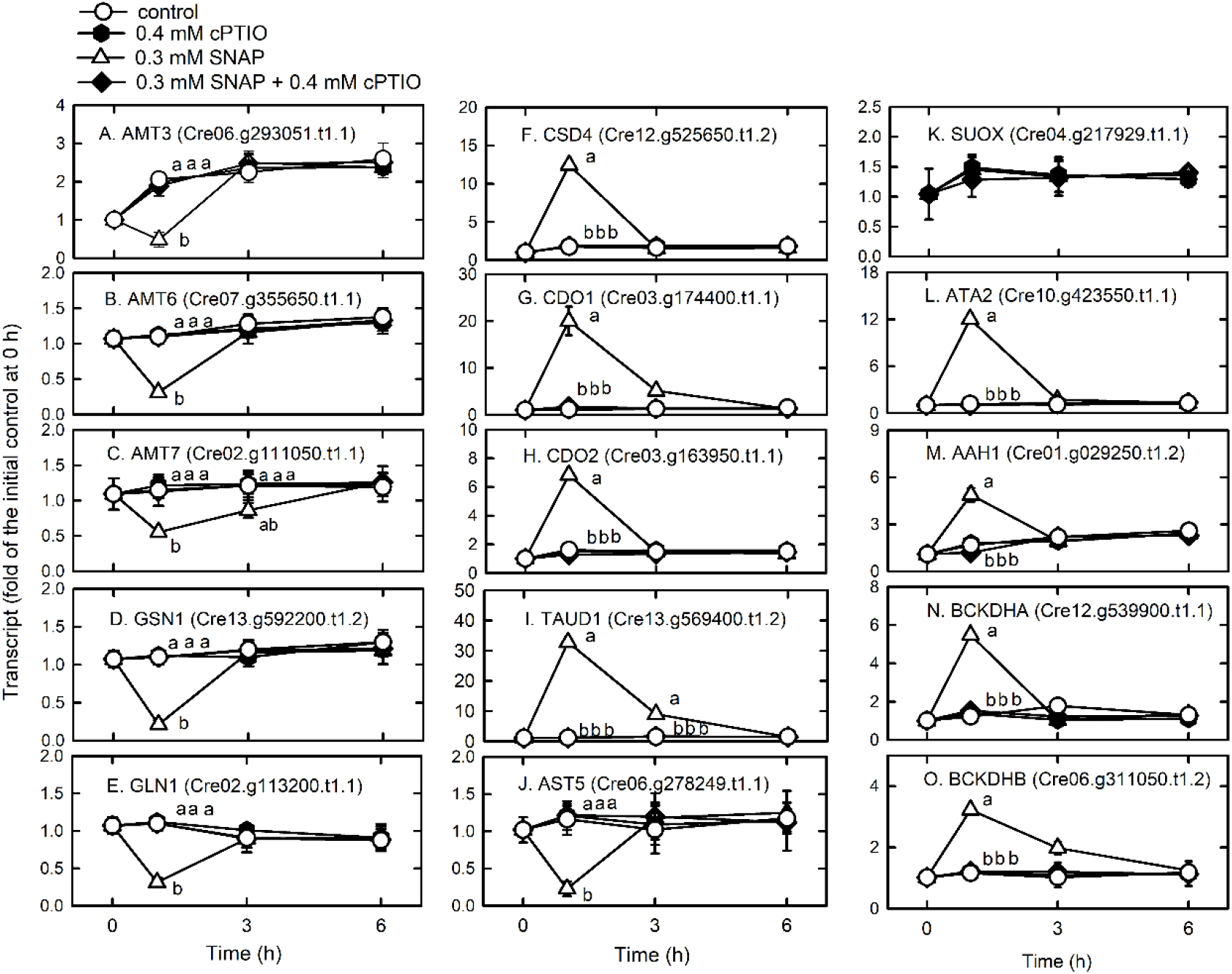
Time-course changes in the transcript abundances of AMT3 (A), AMT6 (B), AMT7 (C), GSN1 (D) GLN1 (E), GDH1 (F), CSD4 (G), CDO1 (H), CDO2 (I), TAUD1 (J), AST5 (K), ATA2 (L), BCKDHA (N), and BCKDHB (O) in *C. reinhardtii* in response to 0.3 mM SNAP in the presence or absence of 0.4 mM cPTIO. Data are expressed as the mean ± SD (n = 3). Different symbols indicate significant differences between treatments (Scheffe’s test, *P* < 0.05).

Cysteine desulfurase (CSD4, Cre12.g525650.t1.2), which moves the sulfur from the cysteine to sulfur-containing recipients, showed an increase in transcriptional expression in the NO treatment (Fig. 5G). Cysteine can be oxidized to 3-sulfinoalanine (Chai et al., 2005) by cysteine dioxygenase (CDO), and the transcript abundances of CDO1 (Cre03.g174400.t1.1) (Fig. 5H) and CDO2 (Cre03.g163950.t1.1) (Fig. 5I) increased after NO treatment. In the NO treatment, the transcript abundance of taurine dioxygenase (TAUD1, Cre13.g569400.t1.2), which degrades taurine to sulfite and aminoacetaldehyde, increased (Fig. 5J) but that of aspartate aminotransferase (AST5, Cre06.g278249.t1.1), which converts 3-sulfinoalanine to pyruvate, decreased (Fig. 5K). Thus, cysteine is catabolized to taurine and then degraded to sulfite instead of pyruvate. Then, sulfite is oxidized by sulfite oxidase (SUOX) to avoid toxicity due to accumulation (Hansch et al., 2007). However, NO did not affect the expression of SUOX (Cre04.g217929.t1.1) (Supplemental Table S3).

The expression of genes related to amino acid degradation was transiently upregulated by NO, including L-allo-threonine aldolase (ATA2, Cre10.g423550.t1.1) (Fig. 5L), the aromatic amino acid hydroxylase-related protein (AAH1, Cre01.g029250.t1.2) (Fig. 5M), and the catabolism of branched-chain amino acids (valine, leucine and isoleucine; 2-oxoisovalerate dehydrogenase E1 component alpha (BCKDHA, Cre12.g539900.t1.1) (Fig. 5N) and beta (BCKDHB, Cre06.g311050.t1.2) (Fig. 5O) (Supplemental Table S6). The presence of cPTIO inhibited the changes induced by SNAP treatment.

NO increased the transcript abundance of periplasmic arylsulfatase (ARS3, Cre10.g430200.t1.2), arylsulfatase (ARS7, Cre01.g011901.t1.1), plasma membrane sodium/sulfate co-transporters (SLT1, Cre12.g502600.t1.2; SLT2, Cre10.g445000.t1.2), the chloroplast sulfate binding protein as the component of chloroplast transporter (SULP3, Cre06.g257000.t1.2), and SNRK2.2 (SAC3, Cre12.g499500.t1.1), an Snf1-like serine/threonine kinase responsible for the repression of expression of genes responsible for sulfur starvation (Davies et al., 1999; Gonzalez-Ballester et al., 2008). NO decreased the transcript abundance of ARS5 (Cre10.g431800.t1.2), ARS11 (Cre04.g226550.t1.2), ARS13 (Cre05.g239750.t1.2), ARS14 (Cre05.g239800.t2.1), and ARS16 (Cre06.g293800.t1.1) (Fig. 6). The presence of cPTIO inhibited these changes in response to SNAP treatment.

**Figure 6.**
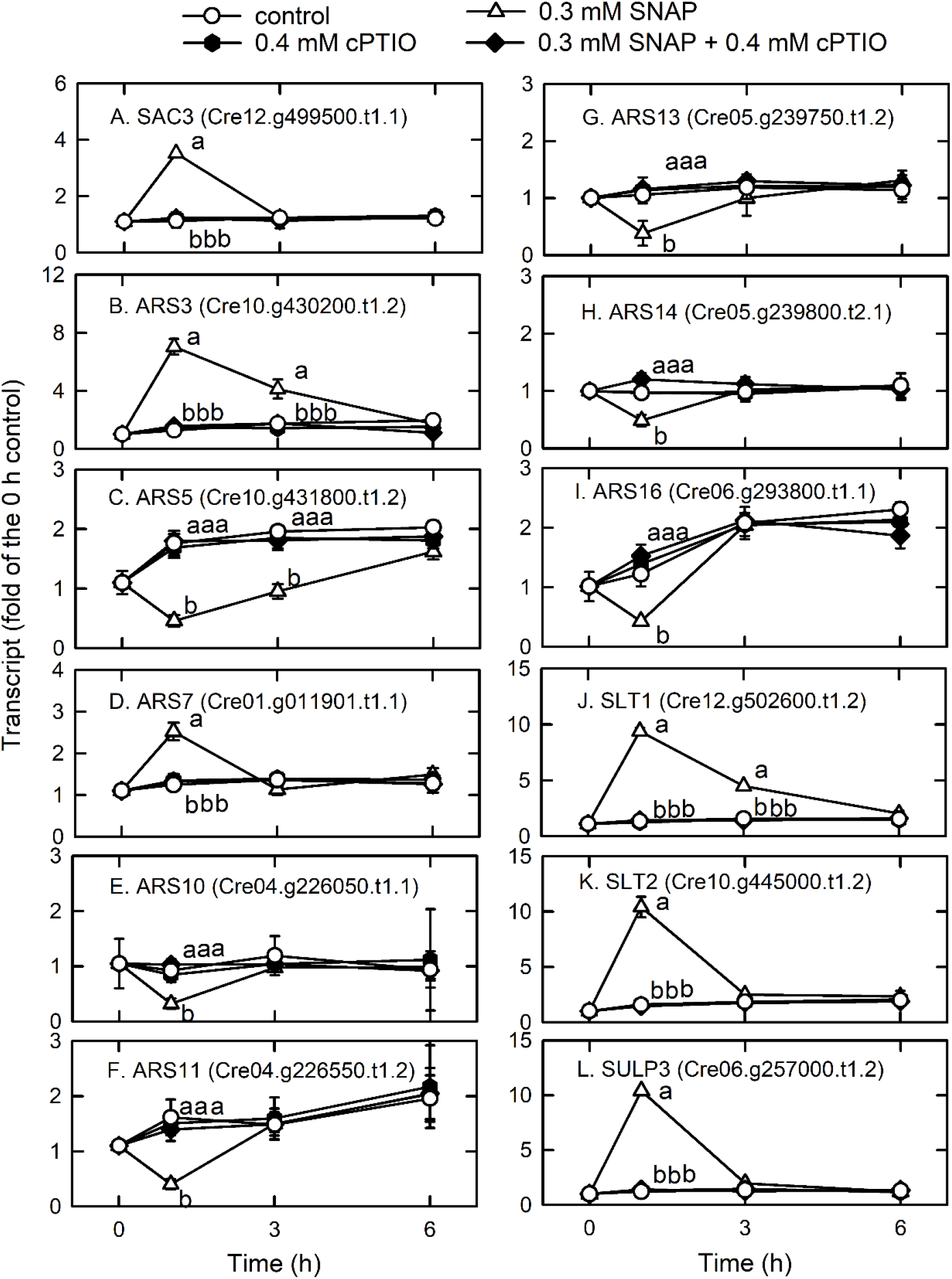
Time-course changes in the transcript abundances of SAC3 (A), SUOX (B), ARS3 (C), ARS7 (E), ARS5 (D), ARS11 (G), ARS13 (H), ARS14 (I), ARS16 (J), SLT1 (K), SLT2 (L), and SULP3 (M) in *C. reinhardtii* after exposure to 0.3 mM SNAP in the presence or absence of 0.4 mM cPTIO. Data are expressed as the mean ± SD (n = 3). Different symbols indicate significant differences between treatments (Scheffe’s test, *P* < 0.05).

### NO Modulates Protein Homeostasis and Quality

Based on the identified GO terms, NO induces protein degradation through ubiquitination and membrane trafficking (Supplemental Fig. S6). For genes involved in the ubiquitin-proteasome system (Supplemental Table S8), the transcript abundances of AAA+-type ATPase (Cre16.g650150.t1.3), which is responsible for the opening of the gates for substrate into the axial entry ports of the proteases (Yedidi et al., 2017); ubiquitin-protein ligase E2 (Cre07.g342506.t1.2); ubiquitin-protein ligase E3 A (UBE3A; Cre08.g364550.t1.3); probable E3 ubiquitin-protein ligase HERC1 (Cre02.g099100.t1.3); and ubiquitin fusion degradation protein (Cre03.g179100.t1.2), which is similar to ubiquitin fusion degradation protein 1, were increased by NO treatment (Fig. 7). In contrast, the transcript abundances of ubiquitin-conjugating enzyme E2I (UBC9, Cre01.g019450.t1.1), which exhibits the activity of small ubiquitin-like modifier (SUMO) E2 conjugase (Wang et al., 2008), a central enzyme in SUMO conjugation for interaction with E1 to accept SUMO in the formation of a SUMO∼UBC9 thioester bond (Pichler et al., 2017), and ubiquitin-conjugating enzyme E2 (UBC3, Cre03.g167000.t1.2) (Fig. 7D) decreased in the NO treatment (Fig. 7). These changes induced by the SNAP treatment were inhibited in the presence of cPTIO.

**Figure 7.**
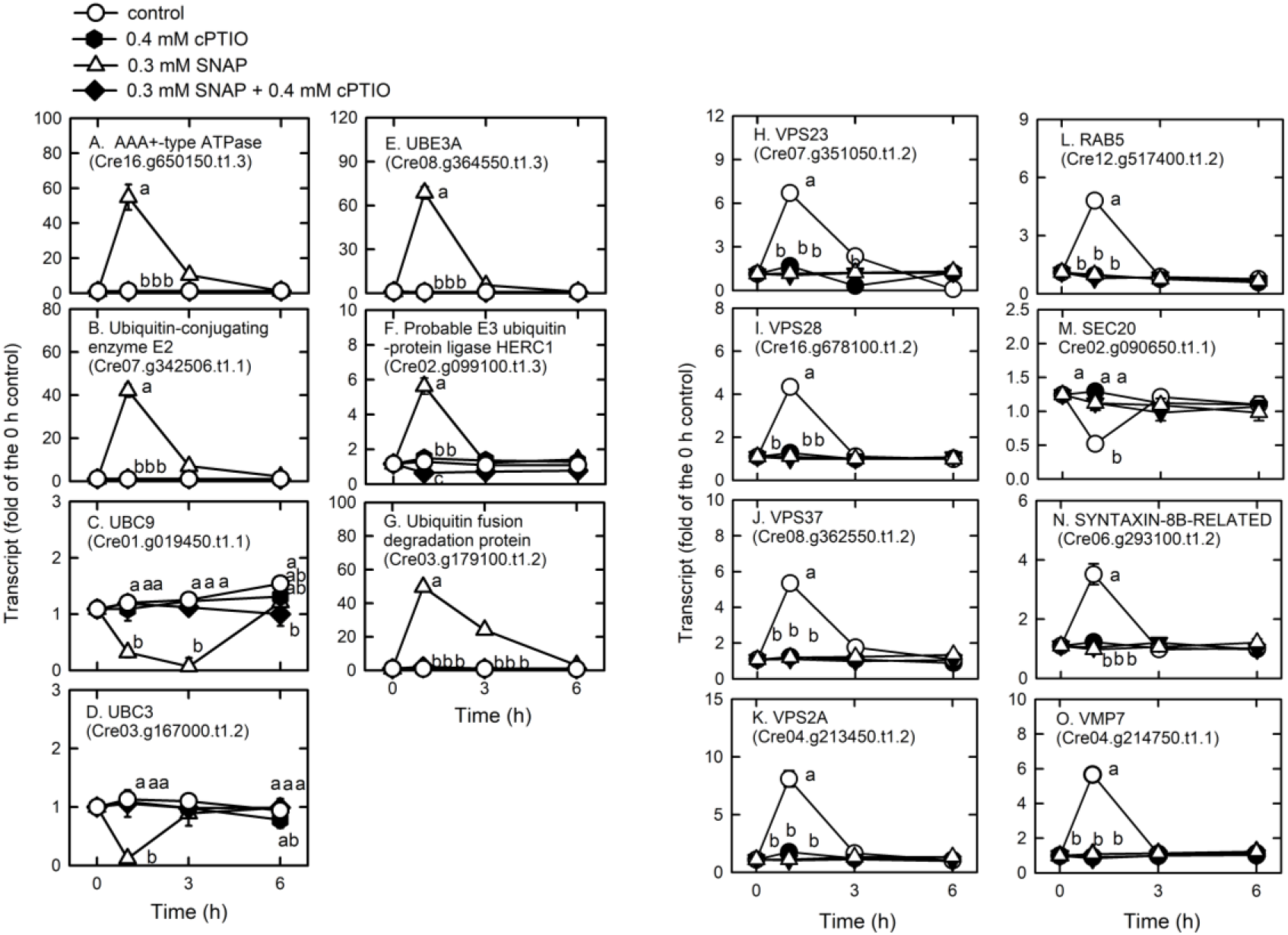
Time-course changes in the transcript abundances of AAA+-type ATPase (A), ubiquitin-protein ligase E2 (B), UBC9 (C), UBC3 (D), UBE3A (E), probable E3 ubiquitin-protein ligase HERC1 (F), ubiquitin fusion degradation protein (G), VPS23 (H), VPS28 (I), VPS37 (J), VPS2A (K), RAB5 (L), SEC20 (M), SYNTAXIN-8B-RELATED (N), and VAMP (O) in *C. reinhardtii* after exposure to 0.3 mM SNAP in the presence or absence of 0.4 mM cPTIO. Data are expressed as the mean ± SD (n = 3). Different symbols indicate significant differences between treatments (Scheffe’s test, *P* < 0.05).

We detected most of the genes encoding endocytosis (Supplemental Table S9A) and SNAREs (Supplemental Table S9B; soluble *N*-ethylmaleimide-sensitive factor attachment protein receptor), which mediate fusion events for autophagosome biogenesis as the membrane trafficking pathways. NO increased the transcript abundances of the vacuolar protein sorting (VPS) proteins, which sort receptors within the endocytic pathway (Bishop and Woodman, 2001) including the subunits of the ESCRT-I (endosomal sorting complex I required for transport) complex (i.e., VPS23 (Cre07.g351050.t1.2), VPS28 (Cre16.g678100.t1.2), and VPS37 (Cre08.g362550.t1.2)), the subunit of the ESCRT-III complex (VPS2A; Cre04.g213450.t1.2), and RAB5 (Cre12.g517400.t1.2), a small rab-related GTPase that regulates vesicle formation and membrane fusion. NO also increased the transcript abundances of SYNTAXIN-8B-RELATED (Cre06.g293100.t1.2), a member of the syntaxin family that is involved in protein trafficking from early to late endosomes via vesicle fusion (Prekeris et al., 1999), and endosomal R-SNARE protein belonging to the VAMP (vesicle associated membrane protein)-like family (R.III) (VMP7, Cre04.g214750.t1.1), which functions in clathrin-independent vesicular transport and membrane fusion events necessary for protein transport from early endosomes to late endosomes (Advani et al., 1999; Prekeris et al., 1999). NO decreased the transcript abundances of endoplasmic reticulum (ER) Qb-SNARE protein and Sec20-family (SEC20, Cre02.g090650.t1.1) (Fig. 7). In addition, NO impacted vesicular protein trafficking that is responsible for the accurate delivery of proteins to correct subcellular compartments (Rosquete et al., 2018) via tethering of a transport vesicle to its target membrane, which is regulated by two classes of tethering factors, long coiled-coil proteins and multi-subunit tethering complexes (MTCs) (Ravikumar et al., 2017). The transport protein particle (TRAPP) complex is a well-studied MTC in yeast and mammals for the regulation of ER-to-Golgi and Golgi-mediated secretion membrane trafficking (TRAnsport Protein Particle, TRAPPI and TRAPPII) and autophagy (TRAPPIII) processes (Kim et al, 2016; Vukasinovic and Zarsky, 2016). Here, we detected several TRAPP genes in *Chlamydomonas*, in which the transcript abundances of TRAPP I subunits (TRS20, Cre13.g571600.t1.2; TRS23, Cre02.g077500.t1.2; TRS31, Cre03.g146467.t1.1; TRS33, Cre16.g663650.t1.2; BET3, Cre03.g145607.t1.1; BET5, Cre06.g289700.t1.2) and a TRAPPIII subunit (TRS85, Cre01.g024400.t1.1) were transiently decreased by NO burst (Fig. 8). As a catabolic process for recycling cellular materials, autophagy is also induced by NO burst (Supplemental Table S9C), which was reflected by a decrease in TOR1 (Cre09.g400553.t1.1) transcript abundance as well as an increase in the transcript abundance of ATG3 (Cre02.g102350.t1.2), ATG4 (Cre12.g510100.t1.1), ATG5 (Cre14.g630907.t1.1), ATG6 (Cre01.g020264.t1.1), ATG7 (Cre03.g165215.t1.1), ATG8 (Cre16.g689650.t1.2), and ATG9 (Cre09.g391500.t1.1) (Fig. 9). However, NO decreased the transcript abundance of ATG12 (Cre12.g557000.t1.2) (Fig. 9I). ATG8 and ATG8-PE (phosphatidylethanolamine) proteins, which were detected using western blot were also increased by NO burst. The changes in the expression of these genes by SNAP were inhibited in the presence of cPTIO (Fig. 9J).

**Figure 8.**
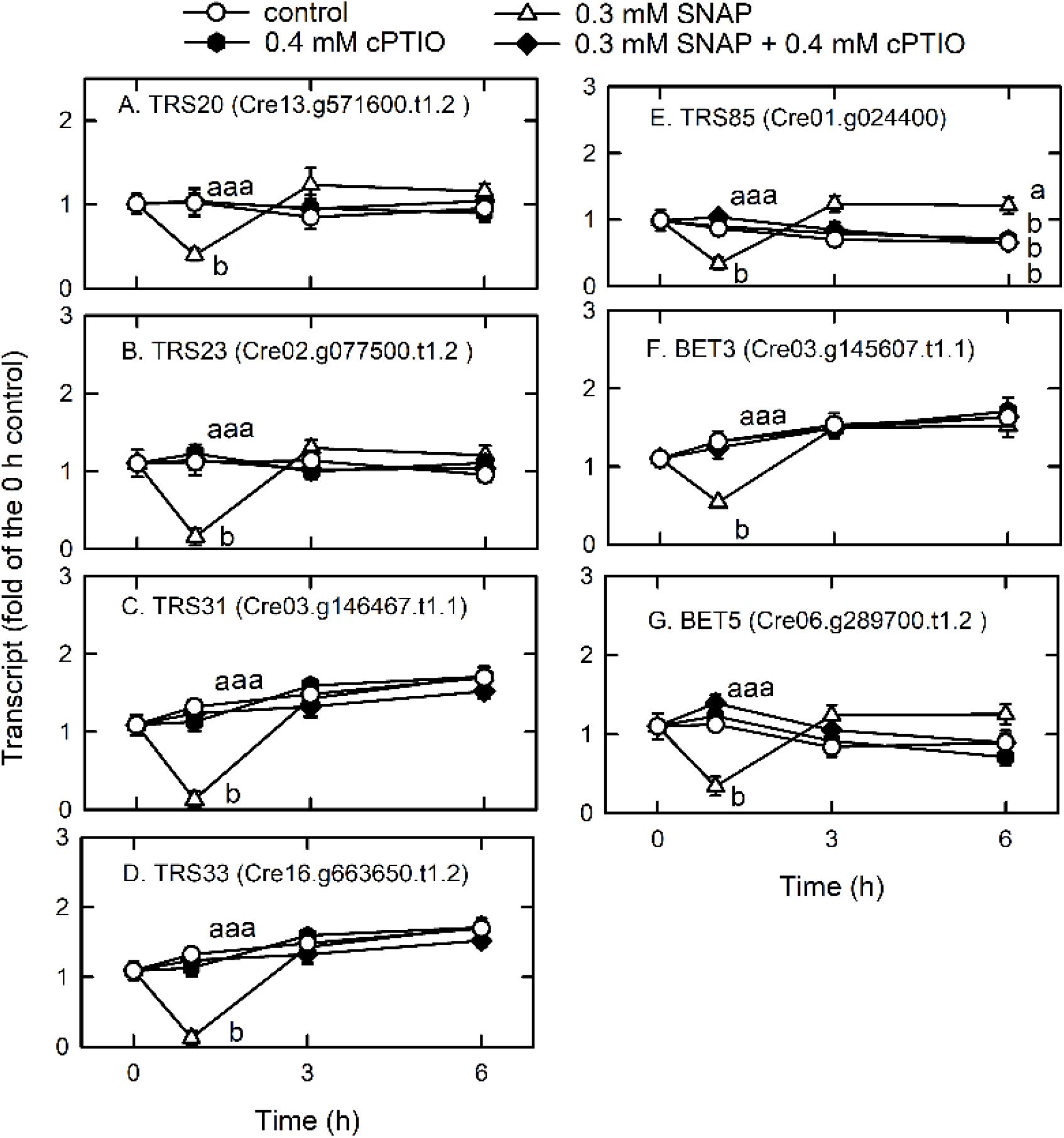
Time-course changes in the transcript abundances of TRS20 (A), TRS23 (B), TRS31 (C), TRS33 (D), TRS85 (E), BET3 (F), and BET5 (G) in *C. reinhardtii* after exposure to 0.3 mM SNAP in the presence or absence of 0.4 mM cPTIO. Data are expressed as the mean ± SD (n = 3). Different symbols indicate significant differences between treatments (Scheffe’s test, *P* < 0.05).

**Figure 9.**
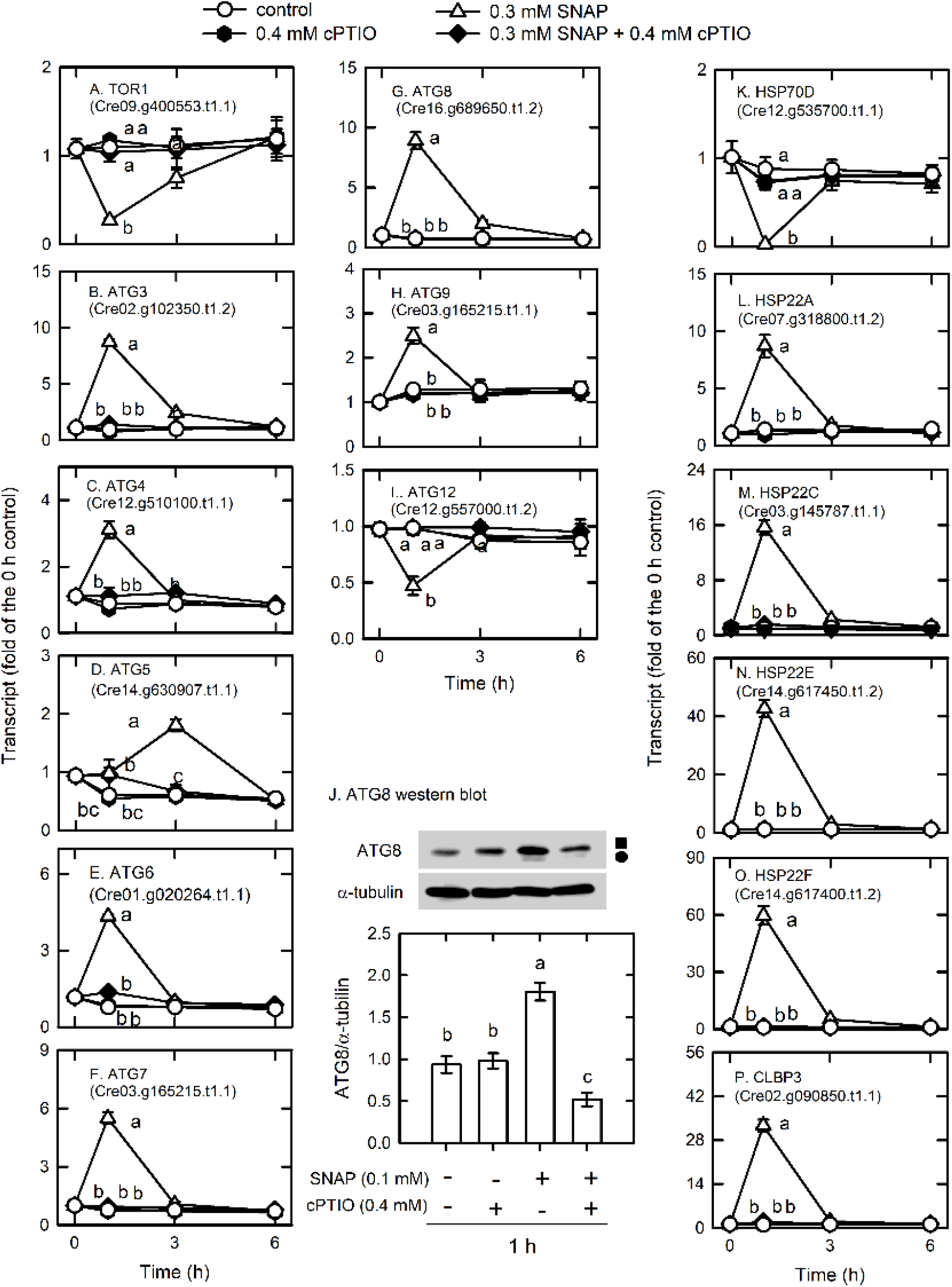
Time-course changes in the transcript abundances of autophagy-related genes and heat-shock proteins and the protein abundances of ATG8 in *C. reinhardtii* upon exposure to 0.3 mM SNAP in the presence or absence of 0.4 mM cPTIO. A. TOR1; B. ATG3; C. ATG4; D. ATG5; E. ATG6; F. ATG7; G. ATG8; H. ATG9; I. ATG12; J. ATG8 (square symbol) and ATG8-PE (circle symbol) protein abundances; K. HSP70D; L. CLBP3; M. HSP22A; N. HSP22C; O. HSP22E; P. HSP22F. Data are expressed as the mean ± SD (n = 3). Different symbols indicate significant differences between treatments (Scheffe’s test, *P* < 0.05).

NO also modulates protein folding via the transcriptional control of heat shock proteins (HSPs) that control protein folding and homeostasis in *Chlamydomonas* (Schroda et al., 2015). An overview of how strongly NO treatment impacts expression levels for each HSP gene, including DNAJ-like proteins, small HSPs, HSP60s, HSP70s, HSP90s, and HSP100s, is provided in Supplemental Table S9D. HSP70D (Cre12.g535700.t1.1) was significantly decreased by NO burst and CLBP3 (Cre02.g090850.t1.1, belonging to the HSP100 family) and small HSPs (HSP22A, Cre07.g318800.t1.2; HSP22C, Cre03.g145787.t1.1; HSP22E, Cre14.g617450.t1.2; HSP22F, Cre14.g617400.t1.2) were significantly increased by NO burst (Fig. 9). The effect of SNAP on the expression of HSPs was inhibited in the presence of cPTIO.

### Induction of the Antioxidant Defense System by NO

Through enzymatic and non-enzymatic routes, the antioxidant defense system is induced by NO treatment (Supplemental Table S10). Superoxide dismutase (SOD), responsible for the dismutaton of O_2_^.−^ to H_2_O_2_, is among the most important antioxidant defense enzymes in plants (Alscher et al., 2002). Six SODs, FSD1 (Cre02.g096150.t1.2), MSD1 (Cre02.g096150.t1.2), MSD2 (Cre13.g605150.t1.2), MSD3 (Cre16.g676150.t1.2), MSD4 (Cre12.g490300.t1.2), and MSD5 (Cre17.g703176.t1.1), were detected to have the most abundant expression for FSD1 (Supplemental Table S10), in which FSD1 and MSD3 transcript abundances were markedly increased by NO treatment (Fig. 10A-F). Furthermore, the ascorbate-glutathione cycle (AGC), an important defense pathway for detoxification of H_2_O_2_, plays a role in the defense of photo-oxidative stress in *Chlamydomonas* (Barth et al., 2014; Lin et al., 2016, 2018; Chang et al., 2017; Yeh et al., 2019; Kuo et al., 2020b) and was also induced by NO. NO increased the transcript abundance of APX1 (Cre02.g087700.t1.2), DHAR1 (Cre10.g456750.t1.2), and GSHR1 (Cre06.g262100.t1.1), but decreased the transcript abundance of APX2 (Cre06.g285150.t1.2), APX4 (Cre05.g233900.t1.2), and MDAR1 (Cre17.g712100.t1.1) (Fig. 10 and Supplemental Fig. S7). The transcript abundance of GSHR2 (Cre09.g396252.t1.1) was not affected by NO (Supplemental Fig. S7). For glutathione (GSH) and ascorbate (AsA) biosynthesis, the transcript abundance of GSH1 (Cre02.g077100.t1.2; Fig. 10J) and VTC2 (GDP-L-galactose phosphorylase, Cre13.g588150.t1.2; Fig. 10K), a key enzyme for AsA biosynthesis (Urzica et al., 2012), increased under NO treatment, while that of GSH2 (Cre17.g708800.t1.1) was not affected (Supplemental Fig. S7E). However, the activity of SOD (Fig. 10L), APX (Fig. 10M), and MDAR (Fig. 10P) was not affected by NO, whereas GR (Fig. 10Q), DHAR (Fig. 10N), and protein (Fig. 10O) did show an increase in activity. These gene expression changes under SNAP treatment were inhibited in the presence of cPTIO (Fig. 10).

**Figure 10.**
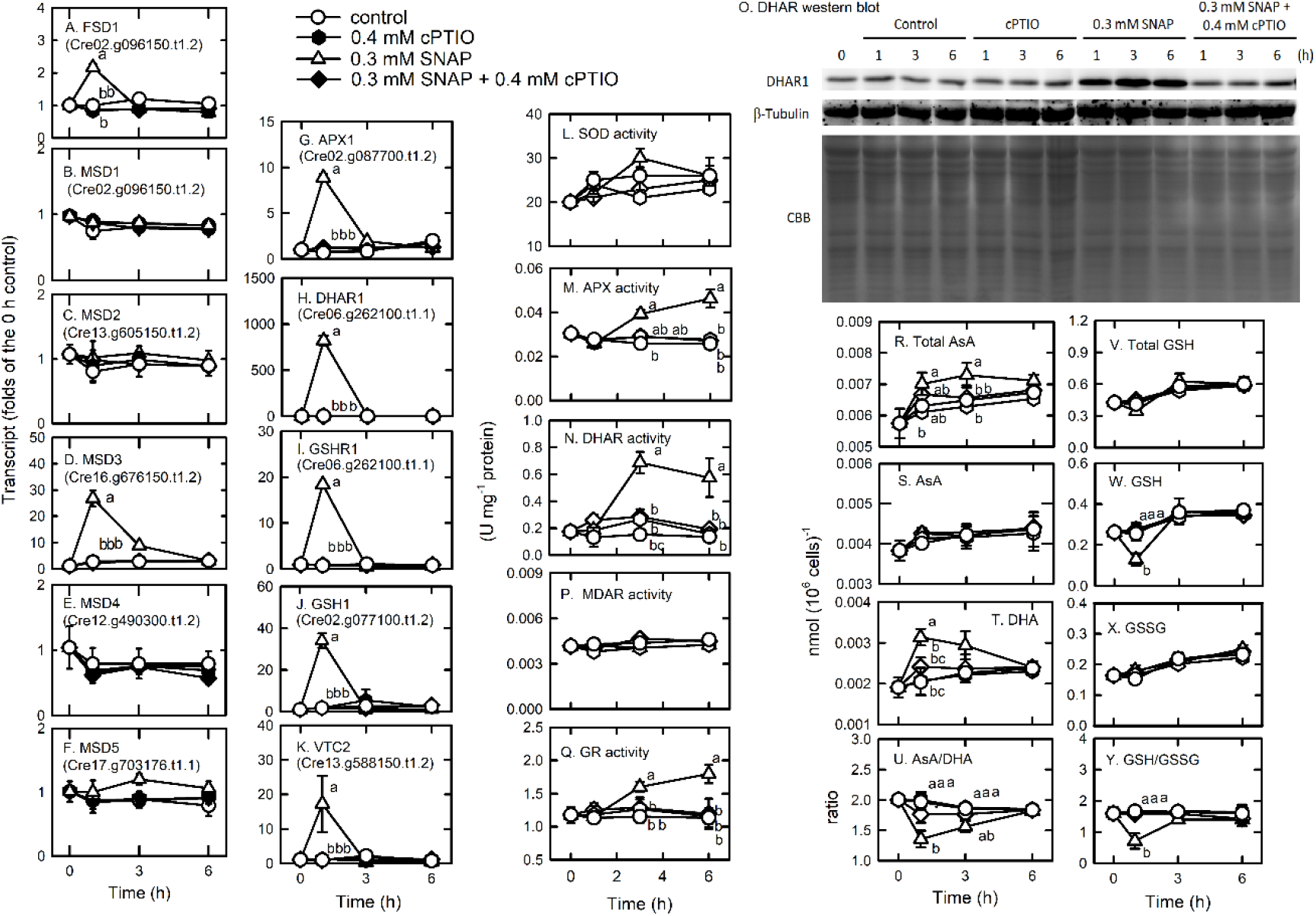
Time-course changes in the transcript abundances of FSD1 (A), MSD1 (B), MSD2 (C), MSD3 (D), MSD4 (E), MSD5 (F), APX1 (G), DHAR1 (H), GSHR1 (I), GSH1 (J), and VTC2 (K); the activity of APX (L), DHAR (M), MDAR (N), and GR (O); the concentrations of total AsA (P), AsA (Q), and DHA (R); the ratio of AsA/DHA (S); the concentrations of total GSH (T), GSH (U), and GSSG (V); and the ratio of GSH/GSSG (W) in *C. reinhardtii* after exposure to 0.3 mM SNAP in the presence or absence of 0.4 mM cPTIO. Data are expressed as the mean ± SD (n = 3). Different symbols indicate significant differences between treatments (Scheffe’s test, *P* < 0.05).

AsA and GSH homeostasis and their redox states (AsA/dehydroascorbate (DHA) and GSH/GSSG (oxidized glutathione)) were affected in the NO treatment. Total AsA (Fig. 10R) and DHA (Fig. 10T) concentrations increased 1 h after NO treatment and AsA concentration was not affected (Fig. 10S). The AsA redox state decreased after 1 h of NO treatment and then recovered (Fig. 10U). Total GSH concentration did not change in the NO treatment (Fig. 10V), while the GSH concentration decreased (Fig. 10W) and the GSSG concentration increased (Fig. 10X), which in turn caused a decrease in the GSH redox state that was restored after 3 h (Fig. 10Y). These changes were inhibited in the presence of cPTIO.

Furthermore, the transcript abundances of a gene homologous to glutathione peroxidase (GPX5, Cre10.g458450.t1.3) and σ-class glutathione-*S*-transferase genes (GSTS1, Cre16.g688550.t1.2; GSTS2, Cre01.g062900.t1.3), associated with oxidative stress acclimation and detoxification response in *Chlamydomonas* (Ledford et al., 2007; Fischer et al., 2012), also increased under NO treatment. GPX1 (Cre02.g078300.t1.1), GPX (Cre08.g358525.t1.1), GPX3 (Cre03.g197750.t1.2), and GPX4 (Cre10.g440850.t1.1) showed a decrease in transcript abundance in the NO treatment (Supplemental Fig. S8).

NO regulates the expression of methionine sulfoxide reductase (MSR), which functions in the reversibility of the oxidization of methionine to methionine and the control of redox homeostasis through modulating the redox status of methionine in proteins (Branlant, 2012; Rey and Tarrago 2018). Here, transcript abundances of MSRA3 (Cre08.g380300.t1.1), MSRA5 (Cre10.g464850.t1.1), and MSRB2.2 (Cre14.g615100.t1.2) increased under NO treatment while those of MSRA2 (Cre06.g257650.t1.2), MSRA4 (Cre01.g012150.t1.2), and MSRB2.1 (Cre14.g615000.t1.1) showed a decrease. The transcript abundance of MSRA1 (Cre13.g570900.t1.1) was not affected by NO treatment (Supplemental Fig. S8). The presence of cPTIO inhibited the changes of MSR gene expression.

## DISCUSSION

NO is a cellular messenger that mediates diverse signaling pathways and plays a role in many physiological processes in plants (Lamattina and Polacco, 2007; Besson-Bard et al., 2008). Studying NO burst over a short-term period provided us a chance to elucidate the metabolic shift to a brief NO attack and the following acclimation processes in *Chlamydomonas* cells. We recently discovered that NO interacts with reactive oxygen species (ROS) to induce cell death in association with autophagy in *Chlamydomonas* cells under high intensity illumination (Kuo et al., 2020a). Fortunately, ROS over-production and oxidative damage were not found in the 0.3 mM SNAP treatment. Thus, the interference of ROS in the short-term response to NO can be excluded. Moreover, the cells collected from the early exponential phase are not nutrient-starved. The role of NO as a factor responsible for SNAP-induced changes was confirmed with the cPTIO treatment, which allowed for a better understanding of the novel components of gene networks in *Chlamydomonas* cells against sub-lethal NO stress using transcriptome and physiological analyses. After 1 h of NO treatment, a decrease in the expression of genes associated with transcriptional and translational activity reflected an inhibition of metabolism in *Chlamydomonas* cells under NO stress. However, *Chlamydomonas* growth was not impacted by sub-lethal NO stress. The present data suggest that the acclimation of *Chlamydomonas* cells to NO burst can be accomplished by specific metabolic pathways.

### Induction of the NO Scavenging System Allows for the Acclimation of *Chlamydomonas* to NO Stress

A transient upregulation of truncated hemoglobin (THB1, Cre14.g615400.t1.2; THB2, Cre14.g615350.t1.2), flavodiiron proteins (FLVB, Cre16.g691800.t1.1), and cytochrome P450 (CYP55B1, Cre01.g007950.t1.1; Supplemental Table S6 and Fig. 12) was found during the period of NO burst. In *Chlamydomonas*, THB1 with dioxygenase activity that converts NO into nitrate (Calatrava et al., 2017) is upregulated by NO (Sanz-Luque et al., 2015b). Sanz-Luque et al. (2015a) showed that treatment with 0.1 mM DEANONOate, a NO donor, increased THB1 transcript abundance in the NIT2 (the nitrate assimilation-specific regulatory gene) wild type and the *nit2* mutant, while THB2 transcript abundance decreased in the NIT2 wild type and showed a slight increase in the *nit2* mutant. *Chlamydomonas* is able to reduce NO to N_2_O via FLVB under light conditions (Chaux et al., 2017) or CYP55B1 in the dark (Burlacot et al., 2019). Accordingly, the present data indicate that the NO scavenging system is induced by NO burst to prevent a toxic NO effect, thus allowing the implementation of acclimation processes post NO exposure.

**Figure 11.**
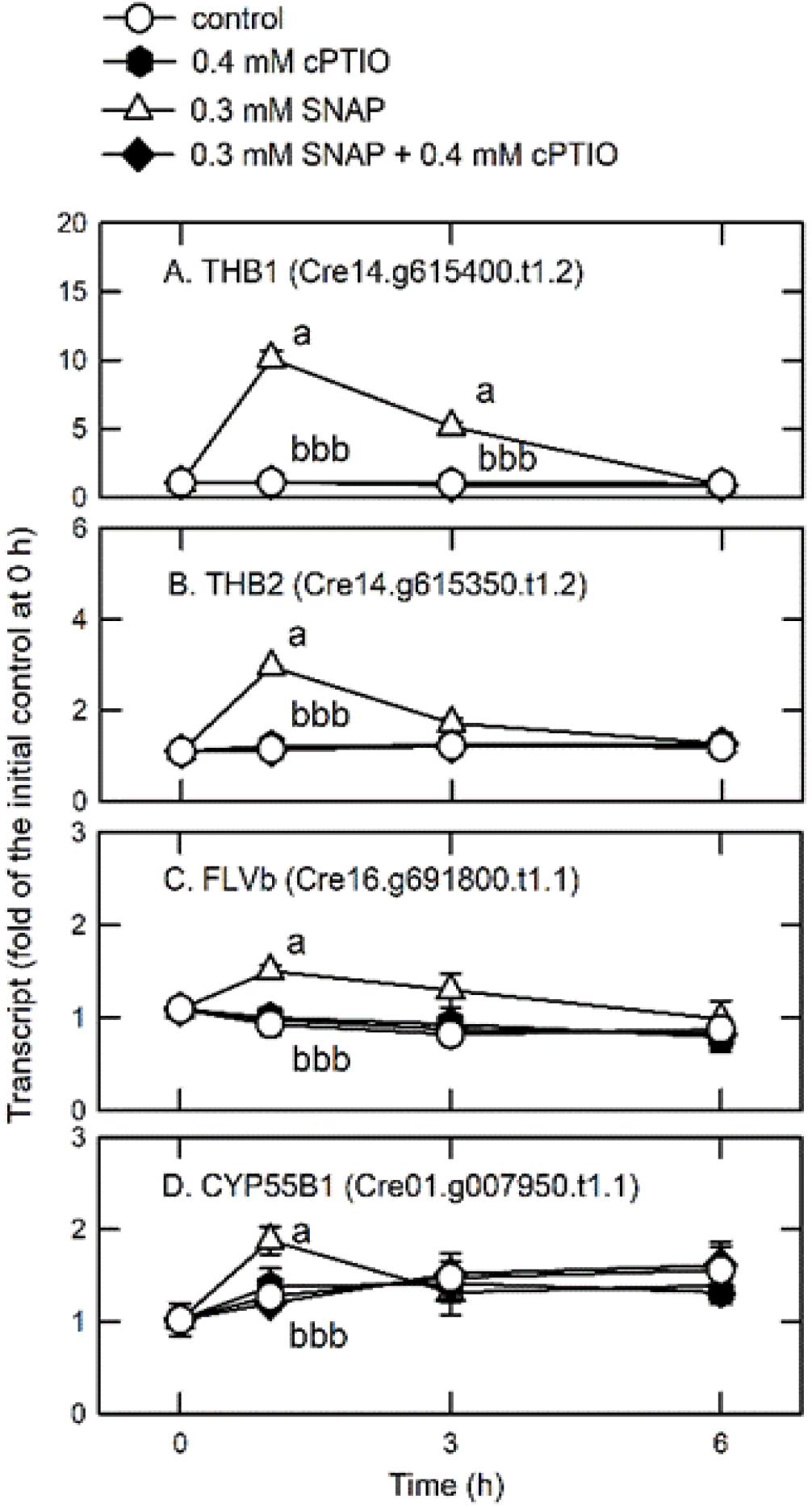
Time-course changes in the transcript abundances of THB1 (A), THB2 (B), FLVB (C), and CYP55B1 (D) in *C. reinhardtii* after exposure to 0.3 mM SNAP in the presence or absence of 0.4 mM cPTIO. Data are expressed as the mean ± SD (n = 3). Different symbols indicate significant differences between treatments (Scheffde’s test, *P* < 0.05).

**Figure 12.**
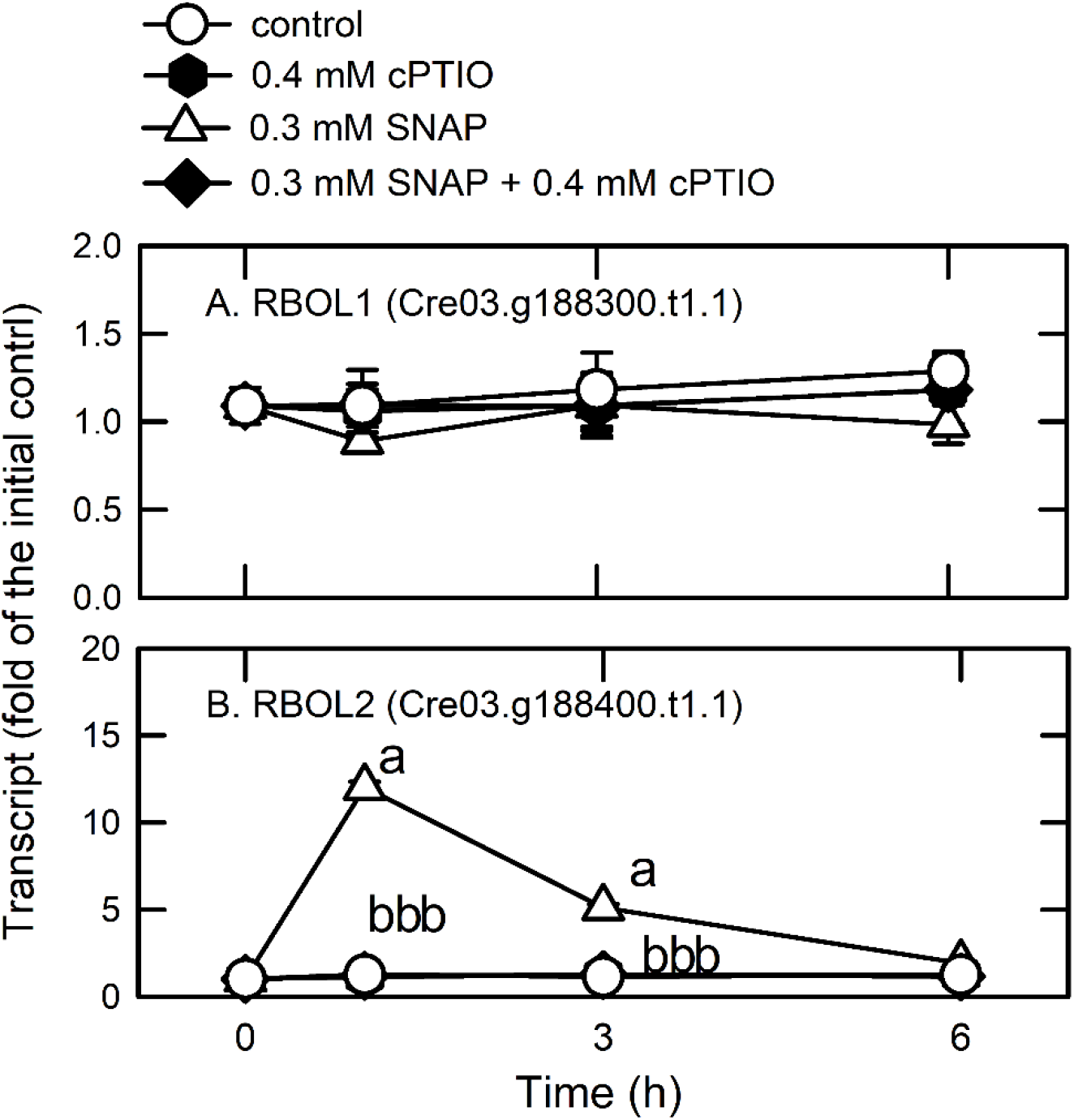
Time-course changes in the transcript abundances of RBOL1 (A) and RBOL2 (B) and the activity of NADPH oxidase (C) in *C. reinhardtii* after exposure to 0.3 mM SNAP in the presence or absence of 0.4 mM cPTIO. Data are expressed as the mean ± SD (n = 3). Different symbols indicate significant differences between treatments (Scheffe’s test, *P* < 0.05).

### NO Alters Nitrogen/Sulfur Assimilation and Primary Amino Acid Biosynthesis

The availability of nitrogen and sulfur are restricted by sudden NO challenge, as reflected by a transient drop in the expression of ammonium transporters and assimilation enzymes, whereas an increased SAC3 expression was observed, which is related to the inhibition of the expression of most ARSs. Moreover, the temporal upregulation of genes involved in the degradation of several amino acids, including sulfur-containing amino acids, under NO stress suggests that NO burst decreased the synthesis of proteins and other nitrogen-containing compounds that use these amino acids as building blocks. NO also causes a reversible inhibition of high-affinity nitrate/nitrite and ammonium transport and NR activity through post-translational regulation (Sanz-Luque et al., 2013). However, time-course changes in transcription levels reveal that the impedance in nitrogen/amino acid and sulfur utilization by NO burst is relieved after 3 h. The upregulation of sulfate transporters (SLT1, SLT2, and SULP3) and GDH1 functioning in the incorporation of ammonium in *Chlamydomonas* (Muñoz-Blanco and Cárdenas, 1989) serves a purpose for improving sulfur availability and amino acid synthesis under NO stress. However, the process by which nitrogen uptake is limited under NO stress is not clear.

### Mechanisms to Overcome NO-Induced Protein Stress

A decrease in the expression of the genes encoding the oligosaccharyltransferase complex, including RNP1 (oligosaccharyltransferase complex subunit delta (ribophorin II)), GTR17 (Glycosyltransferase), GTR22, GTR25 (oligsaccharyltransferase STT3 subunit), STT3B (dolichyl-diphosphooligosaccharide-protein glycosyltransferase subunit STT3B), CANX (calnexin (CANX)), and CRT2 (Calreticulin 2, calcium-binding protein), by NO treatment (Supplemental Table S6) reflects the dysfunction of proteins caused by a blockage of glycosylation in the ER. However, the expression of DAD1, which functions in N-linked glycosylation in the ER (Kelleher and Gilmore, 1994), increased in the NO treatment. The NO-induced downregulation of TRAPPI and III proteins, which are related to the transport of proteins to correct compartments and the formation of autophagosomes (Garcia et al., 2019; Rosquete et al., 2019), also suggests that there was transient transfer of misfolded and damage proteins during NO treatment. Furthermore, the downregulation of SUMO E2 conjugase, UBC9, by NO reflects the inhibition of SUMOylation for post-translational modification of the proteins, which is crucial for the growth of *Chlamydomonas* cells (Knobbe et al., 2015). Clearly, NO burst causes protein stress in *Chlamydomonas*.

To overcome disorders caused by potentially misfolded proteins, the membrane trafficking system is induced for the degradation of damaged proteins in the lysosome and/or vacuole via autophagy, which was reflected by a transient increase in the expression of E2 and E3 ubiquitin ligases, ESCRT subunits (VSP), SNAREs, and ATGs by NO treatment. The increase in the expression of SYNTAXIN-8B-RELATED and VAMP7, which are endosomal syntaxins that mediate the steps of endosomal protein trafficking (Prekeris et al., 1999), in *Chlamydomonas* cells suggests that there was fusion between autophagosomes (late endosomes) and lysosomes under NO stress. The TEM images show the formation and fusion of large vesicles at 1 h, possibly acting to enclose the cytosolic components for degradation (Xie and Klionsky, 2007; Nakatogawa et al., 2009); the large vesicles were not visible after 6 h (Supplemental Fig. S9). Using autophagy as a lysosome-mediated pathway for the degradation of cytosolic proteins and organelles (Choi et al., 2013), ATG7 is involved in the two ubiquitin-like systems necessary for the selective cytoplasm-to-vacuole targeting (Cvt) pathway and autophagy by activation of ATG12 and ATG8 and assigns them to E2 enzymes, ATG10 and ATG3, respectively. During this process, ATG8 conjugates PE to form lipidated ATG8 (ATG8-PE), an autophagosome membrane component and binding partner for autophagy receptors, which act to recruit cargo to lysosomes for catabolism in ubiquitin-dependent and independent manners for selective autophagy. Although ATG12 and VTC1 (vacuolar transport chaperone-like protein, Cre12.g510250 t1.2) were downregulated by NO burst, an increase in the expression of other ATG genes and abundances of ATG/ATG8-PE proteins in NO-treated *Chlamydomonas* cells demonstrates the induction of autophagy by working with the sorting protein factors involved in the membrane trafficking system for protein degradation and recycling under NO stress. Because the inhibition of TOR activity can trigger autophagy, as evidenced by the increased ATG8 and ATG8-PE proteins in *Chlamydomonas* (Pérez-Pérez et al., 2010), the decreased expression of TOR1 due to NO treatment indicates that TOR plays a role in autophagy induction. In addition to membrane trafficking, protein chaperone systems (HSP22A, HSP22C, HSP22E, HSP22F, and CLBP6) are induced in *Chlamydomonas* cells under NO stress. Thus, the protein trafficking system (ubiquitination, SNARE, autophagy) and HSPs are evoked as a way to remove aberrantly folded or damaged proteins, allowing *Chlamydomonas* cells to maintain normal functions under NO stress.

### NO Inhibits Photosynthesis but Induces the Antioxidant Defense System for the Prevention of Oxidative Stress upon Exposure to NO Burst

A decrease in the expression of genes encoding proteins related to photosynthesis and photosynthetic activity in *Chlamydomonas* by NO illustrates the NO-mediated downregulation of PSII activity and the evolutionary rate of photosynthetic O_2_ at the transcriptional level. Because of the suppression of PSII activity and the photosynthetic O_2_ evolution rate by the NO treatment at the protein level in higher plants (Takahashi and Yamasaki, 2002; Singh et al., 2007; Wodala et al., 2008; Ördög et al., 2013), the effects of NO on photosynthesis by protein modification cannot be ignored. NO is also a factor leading to the degradation of the cytochrome *b*_6_*f* complex and Rubisco through FtsH and Clp proteases in sulfur (de Mia et al., 2019) or nitrogen (Wei et al., 2014) starved *Chlamydomonas* cells. However, it is not clear whether these two protein complexes are degraded under NO burst in nutrient sufficient conditions. We did find that the transcript abundances of the FtsH-like proteases FHL4 (Cre13.g568400.t1.2) and FHL6 (Cre03.g201100.t1.2), and DEG11 (Cre12.g498500.t1.2) were increased by NO burst, whereas the transcript abundances of the ClpP proteases ClpP4 (Cre12.g500950.t1.2) and ClpP5 (Cre12.g486100.t1.2), and the non-catalytic subunit of the ClpP complex, ClpR2 (Cre16.g682900.t1.2) decreased (Supplemental Table S6). However, the role of FtsH-like and DEG proteases responsible for chloroplast protein degradation by NO burst need to be identified. Because the inhibition of photosynthesis is considered the mechanism for algae to acclimate to nitrogen or sulfur limitation to avoid photo-damage (Peltier and Schmidt, 1991; Grossman, 2000; Saloman et al., 2012), our findings suggest that the transient inhibition of photosynthesis can prevent *Chlamydomonas* cells from over-producing ROS, which may occur during NO burst. ROS scavenging ability was enhanced with the increased concentration of AsA due to the upregulation of VTC2 and its regeneration rate, owing to the enhanced DHAR expression (transcript abundance, protein, enzyme activity). In addition, the enhanced GSH regeneration rate, supported by the increase in the GR activity and GSHR1 expression as well as increased GSTS1, GSTS2, and GPX5 expression, prevented the over-production of ROS. Tocopherols (Havaur et al., 2005; Krieger-Liszkay and Trebst, 2006) and carotenoids (Ramel et al., 2012) are also ^1^O_2_ quenchers, but their concentrations and the transcript abundances of their biosynthetic enzymes decreased in the NO treatment (Supplemental Fig. S10). Instead, an increase in the concentration of AsA, an ^1^O_2_ quencher (Kramarenko et al., 2006), was responsible for ^1^O_2_ scavenging under NO stress. NO also triggers MSR expression. Although the redox of AsA and GSH as well as the thiol-based redox regulation proteins, TRX2 (thioredoxin-like protein, Cre03.g157800 t1.1), PRX1 (2-cys peroxiredoxin, chloroplastic, Cre06.g257601 t1.2), and PRX2 (2-cys peroxiredoxin, Cre02.g114600 t1.2; Supplemental Table S6), were initially decreased by NO burst, they recovered 3 h after treatment (the restoration of TRX2, PRX1, and PRX2 transcript abundance is not shown). Thus, the antioxidant defense system and MSR worked together to prevent oxidative stress and the abrupt loss in cellular redox homeostasis occurring under NO stress.

### Burst Oxidase-Like 2-Dependent Regulation of the Expression of Selective NO-Inducible Genes for Coping with NO Stress

NADPH oxidase acts as an important molecular hub during ROS-mediated signaling in plants (Marino et al., 2011) and in the interplay of ROS and NO signalling pathways for the regulation of plant metabolism (Suzuki et al., 2011). A previous study has shown that NADPH oxidase is regulated by transcriptional and enzyme activity levels in higher plants (Hu et al., 2020). The NO-mediated suppression of NADPH oxidase activity is due to post-translational modification known as *S*-nitrosylation (Wang et al., 2013). The analysis of cis-regulatory elements in the gene promoter region of NADPH oxidase showed diverse expression patterns under different conditions (Wang et al., 2013; Kaur and Pati, 2016). In this study, we did not examine whether the *S*-nitrosylation of NADPH oxidase occurs in *Chlamydomonas* upon exposure to NO; however, our present data did detect the constant expression of the gene encoding respiratory burst oxidase-like 1 (RBOL1, Cre03.g188300.t1.1), an NADPH oxidase gene (Fig. 12A). In addition, we observed the significant upregulation of RBOL2 (Cre03.g188400.t1.1) (Fig. 12B) in *Chlamydomonas* after 1 h of NO exposure, followed by the subsequent restoration to a baseline level after 3 h. The activity of NADPH oxidase also increased after exposure to NO (Fig. 12C). Therefore, it is interesting for us to examine whether NADPH oxidase mediates the NO-induced changes in the expression of genes in *Chlamydomonas* using the *rbol2* mutant (Fig. 13A,B). Compared to the wild type, the *rbol2* mutant had a 66% decrease in NADPH oxidase activity with a low RBOL2 transcript abundance (Fig. 13C), indicating a differential regulation pattern on the expression of the genes upon exposure to NO (0.3 mM SNAP). In response to NO treatment (0.3 mM SNAP), the transcript abundance of RBOL1 (Fig. 13Z) was not affected in the *rbol2* mutant and the induction of RBOL2 expression in wild type (CC-5325) was not observed in the *rbol2* mutant (Fig. 13AB). In addition, the NO-induced increase in the expression of THB2 (Fig. 13D), CDO1 (Fig. 13E), CDO2 (Fig. 13F), HSP22A (Fig. 13G), HSP22C (Fig. 13H), ATG4 (Fig. 13J), ATG8 (Fig. 13K), LHCBM9 (Fig. 13L), GUN4 (Fig. 13M), APX1 (Fig. 13P), GSHR1 (Fig. 13R), GSTS1 (Fig. 13T), GSTS2 (Fig. 13U), DHAR1 (Fig. 13W), and PDS1 (Fig. 13Y) in the wild type was not observed in the *rbol2* mutant. In contrast to the NO-mediated inhibition of TOR1 (Fig. 13I) and CYC4 (Fig. 13N) expression in the wild type, the expression of TOR1 and CYC4 showed a significant increase in the *rbol2* mutant after NO treatment. The expression pattern of RBCS1 (Fig. 13P), VTC2 (Fig. 13R), GSH1 (Fig. 13S), MDAR1 (Fig. 13V), and MDAR5 (Fig. 13X) in the *rbol2* mutant in the NO treatment was similar to that in the wild type. Compared to the wild type, the NADPH oxidase activity in the *rbol2* mutant did not increase after SNAP treatment (Fig. 14A). The *rbol2* mutant showed a higher sensitivity to NO stress than the wild type, as reflected by a decrease in *rbol2* mutant viability (Fig. 14B) accompanied with bleaching (Fig. 14C) as the SNAP concentration increased ≥ 0.3 mM. Overall, NADPH oxidase (RBOL2) is associated with the expression of the genes encoding proteins for NO scavenging, antioxidant defense system, autophagy, and heat shock proteins that help to resist NO stress in *Chlamydomonas*.

**Figure 13.**
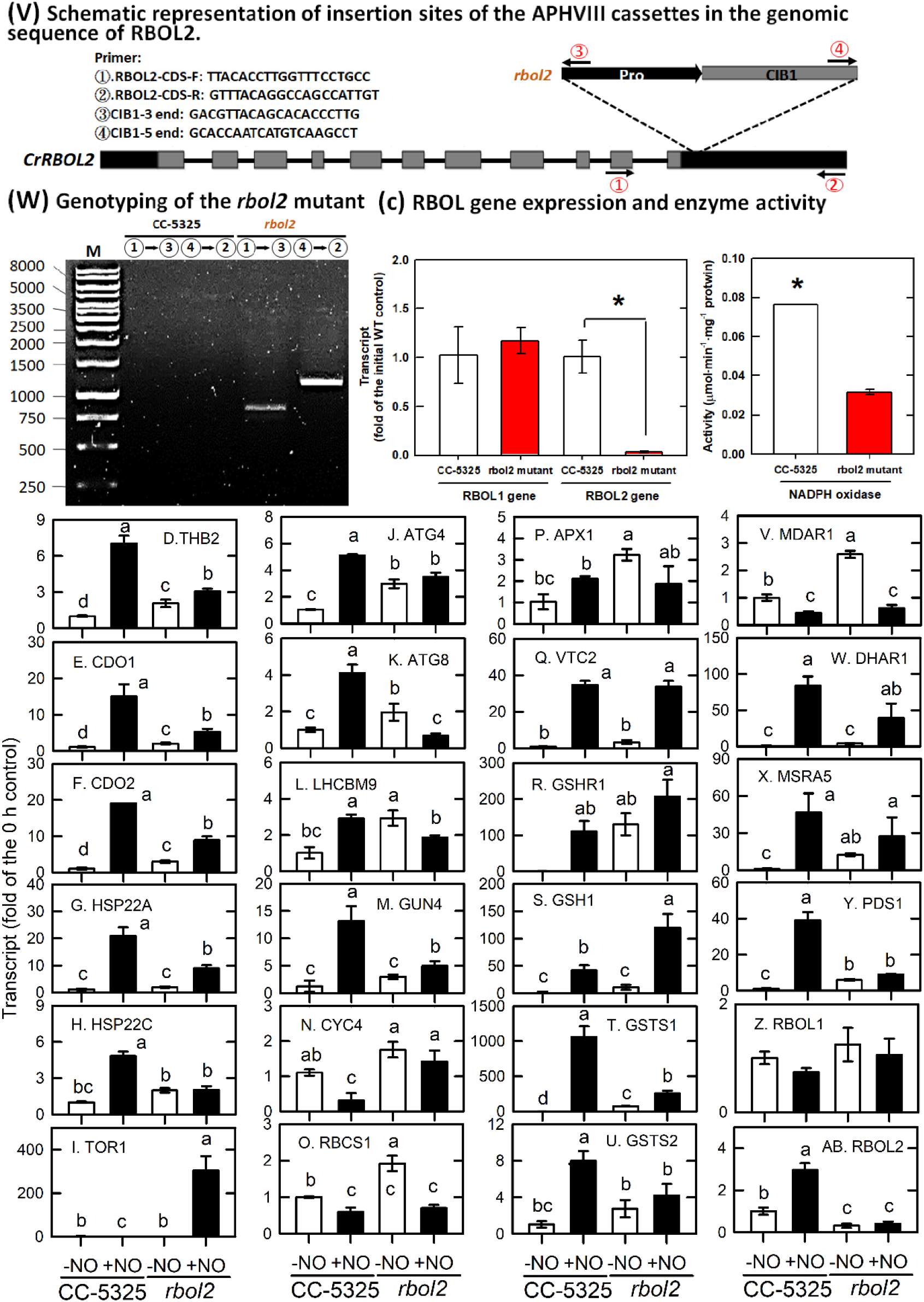
Isolation of the *Chlamydomonas* rbol2 mutant and its transcript abundances of several genes in response to 0.3 mM SNAP in the presence or absence of 0.4 mM cPTIO. (A) Schematic representation of insertion sites of the APHVIII cassettes in the genomic sequence of RBOL2. Tall boxes denote exons, gray boxes indicate protein coding regions and filled boxes show 5’ and 3’ untranslated regions (UTRs) and the promoter region. Arrows indicate primer locations used to detect APHVIII cassette insertions. (B) Genotyping of the rbol2 mutant. Genomic DNA fragments were amplified by PCR using the primer sets indicated in (A). (C) The rbol2 mutant showed low RBOL2 transcript abundance and NADPH oxidase activity but a normal RBOL1 expression compared to the CC-5325 wild type. The data are expressed as the mean ± SD (n = 3) from three independent biological replicates, and * indicates a significant difference between CC-5325 and the rbol2 mutant using *t*-test (*P* < 0.05). The transcript abundance of THB2 (D), CDO1 (E), CDO2 (F), HSP22A (G), HSP22C (H), TOR1 (I), ATG4 (J), ATG8 (K), LHCBM9 (L), GUN4 (M), CYC4 (N), RBCS1 (O), APX1 (P), VTC2 (Q), GSHR1 (R), GSH1 (S), GSTS1 (T), GSTS2 (U), MDAR1 (V), DHAR1 (W), MSAR5 (X), PDS1 (Y), RBOL1 (Z), and RBOL2 (AB). The data are expressed as the mean ± SD (n = 3), in which different symbols indicate significant differences between treatments (Scheffe’s test, *P* < 0.05).

**Figure 14.**
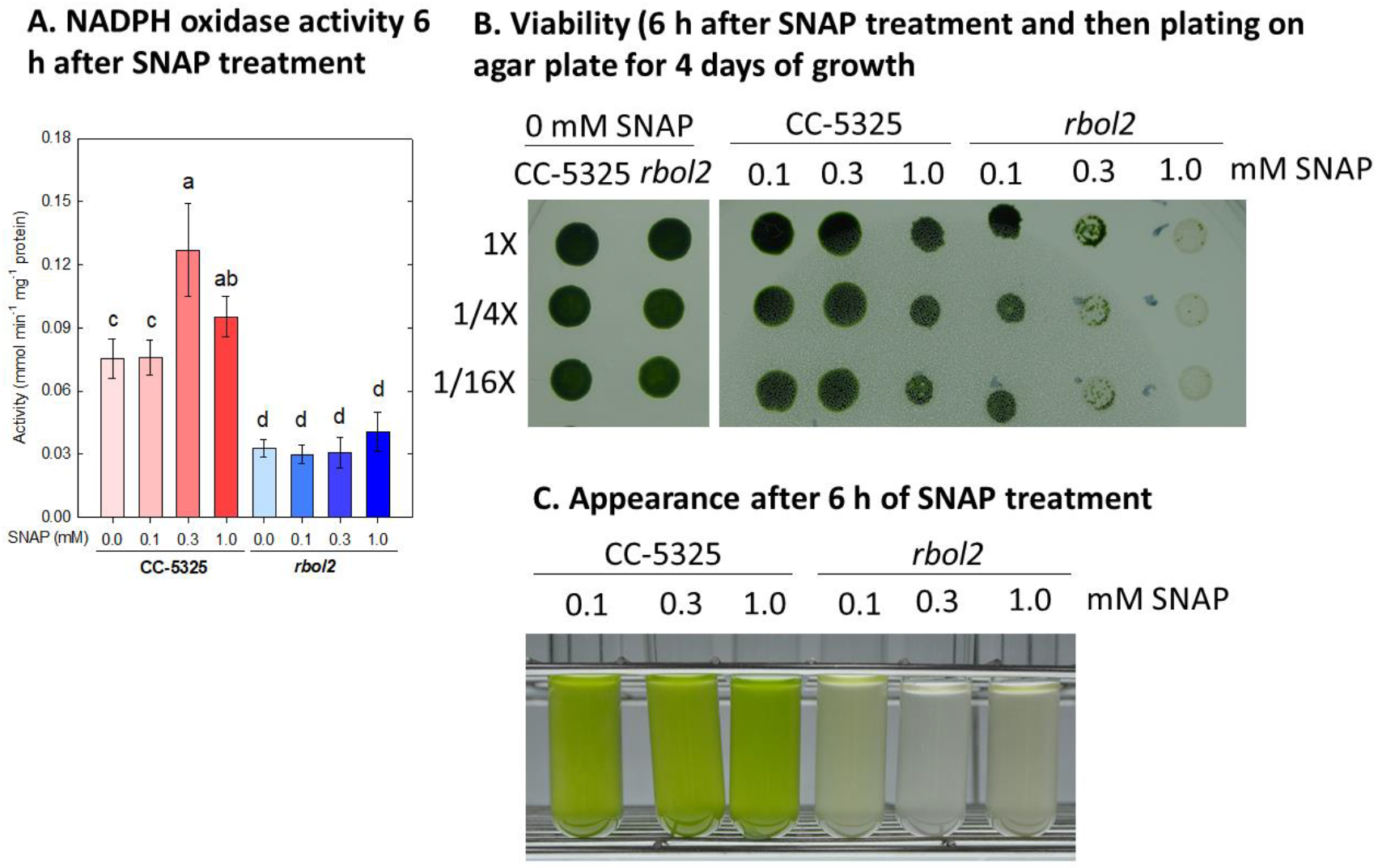
The effects of SNAP treatment on NADPH oxidase activity (A), the viability of *Chlamydomonas* after 4 days of culture on an agar plate including 6 h SNAP treatment (B) and the appearance of cells 6 h after SNAP treatment. SNAP was applied at 0, 0.1, 0.3, or 1.0 mM concentrations. Data are expressed as the mean ± SD (n = 3) for NADPH oxidase activity. Different symbols indicate significant differences between treatments (Scheffe’s test, *P* < 0.05).

## CONCLUSIONS

The acclimation of *Chlamydomonas* to sub-lethal NO stress comprises a temporally orchestrated implementation of metabolic processes including 1. a decrease in the NO amount due to scavenging elements, 2. acclimation to nitrogen and sulfur starvation, 3. upregulation of the protein trafficking system (ubiquitin, SNARE, and autophagy) and molecular chaperone system for dynamic regulation of protein homeostasis and function, and 4. transient inhibition of photosynthesis together with the induction of the antioxidant defense system and modification of the redox state to prevent oxidative stress. NADPH oxidase (RBOL2)-dependent molecular events underlie part of the acclimation mechanisms in *Chlamydomonas* related to coping with NO stress. The direct targets of NO and the mechanisms for the transcriptional regulation of RBOL2 in *Chlamydomonas* are needed to be clarified in the future.

## MATERIALS AND METHODS

### Algal Culture and SNAP Treatments

The green alga *Chlamydomonas reinhardtii*, strain CC-125 (mt-), was obtained from the Chlamydomonas Resource Center (USA) and photoheterotrophically cultured in Tris-acetate phosphate medium (TAP) (Harris, 1989) with a trace element solution in 125 mL flasks (PYREX, Germany) and agitated on an orbital shaking incubator (model OS701, TKS company, Taipei, Taiwan) (150 rpm) under continuous illumination with white light (50 μmol·m^−2^·s^−1^) at 25°C. For chemical treatments, 50 mL cultures were grown to a cell density of 3–5×10^6^ cells·mL^−1^, and after centrifugation at 1,600 ×*g* for 3 min, the supernatant was discarded. The pellet was suspended in fresh TAP medium and centrifuged again. Then, the pellet was re-suspended in fresh TAP medium to a cell density of 3×10^6^ cells·mL^−1^. Ten millilitres of culture was transferred to a 100-mL beaker (internal diameter: 3.5 cm) for pre-incubation at 25°C in 50 μmol·m^−2^·s^−1^ conditions for 1.5 h in an orbital shaker (model OS701, TKS company, Taipei, Taiwan) at a speed of 150 rpm. Then, the algal cells were subjected to treatments at 25°C. SNAP was treated in different concentrations from 0.1, 0.3, to 1.0 mM in the presence or absence of 0.4 mM cPTIO. DMSO was used as the control because SNAP was dissolved in DMSO. Each treatment included three biological independent replicates (n=3). For the determination of cell growth and the estimation of several physiological and biochemical parameters, the number of cells in a 1-mL sample was counted using a hemocytometer. Samples taken before (0 min) and after treatment were centrifuged at 5,000 ×*g* for 5 min, and the pellet was fixed in liquid nitrogen and stored in a −-70°C freezer until analysis.

### Detection of NO Production

An NO-sensitive fluorescent dye, DAF-FM diacetate (Invitrogen Life Technologies, Carlsbad, CA, USA) (Kojima et al., 1998), was used to measure NO production following our previous studies. DAF-FM diacetate is a pH-insensitive fluorescent dye that emits fluorescence after reaction with an active intermediate of NO (Kojima et al., 1998; Chang et al., 2013). The cells were pre-incubated in TAP medium containing 5 μM DAF-FM diacetate for 60 min at 25°C under 50 μmol·m^−2^·s^−1^ conditions, then washed twice with fresh TAP medium, and transferred to 50 μmol·m^−2^·s^−1^ for chemical treatment. The fluorescence was detected via fluorescence microscopy and spectrophotometry.

### Determination of Chlorophyll *a* Fluorescence

Chlorophyll *a* fluorescence parameters were employed to determine the activity of PSII using an AP-C 100 (AquaPen, Photon Systems Instruments, Brno, Czech Republic). A 0.5-mL aliquot of an algal culture was diluted with TAP medium to an OD_750_ = 0.10–0.15 with a chlorophyll *a* content of 1.5–2.2 μg·mL^−1^. A 2-mL aliquot of diluted algal cells was then transferred to an AquaPen cuvette and subjected to a pulse of saturating light of 4,000 μmol·m^−2^·s^−1^ PAR to obtain the light-adapted minimal fluorescence (*F*_t_) and the light-adapted maximal fluorescence (*F*_m_’). To determine the maximum PSII activity, *F*_v_/*F*_m_ (=*F*_m_ - *F*_o_/*F*_m_), 2 mL of diluted algal cells in an AquaPen cuvette was incubated in the dark for 20 min and then flushed with saturated light (4,000 μmol photons·m^−2^·s^−1^) to obtain the dark-adapted minimal fluorescence (*F*_o_) and dark-adapted maximal fluorescence (*F*_m_). The active PSII activity, *F*_v_’/*F*_m_’ = *F*_m_’ - *F*_t_/*F*_m_’, and the maximum PSII activity, *F*_v_/*F*_m_ = *F*_m_– *F*_o_/*F*_m_, were then calculated.

The rapid induction of chlorophyll *a* fluorescence was also determined via the OJIP test (Strasser and Strasser, 1995; Strasser et al., 2000). The inflections in the O-J-I-P curve represent the heterogeneity of the process during photochemical action; peak J represents the momentary maximum level of Q_A_, Q_A_ Q_B_, and Q_A_ Q_B_ ; peak I represents the level of Q_A_ Q_B_ ; and peak P represents the maximum level of Q_A_, Q_B_, and PQH_2_ (Stirbet et al., 1998). The fluorescence values at time intervals corresponding to the O-J-I-P peaks were recorded as follows: *F*_o_ = fluorescence intensity at 50 µs; *F*_J_ = fluorescence intensity at peak J (at 2 ms); *F*_i_ = fluorescence intensity at peak I (at 60 ms); *F*_m_ = maximal fluorescence intensity at the peak P; *F*_v_ = *F*_m_ - *F*_o_ (maximal variable fluorescence); V_j_ = (*F*_j_ - *F*_o_)/(*F*_m_ - *F*_o_); V_i_ = (*F*_i_ - *F*_0_)/(*F*_m_ - *F*_o_); and *F*_m_/*F*_o_; *F*_v_/*F*_o_; *F*_v_/*F*_m_.

### Enzyme Assay and Determination of AsA and GSH

SOD activity was assayed according to (Yashida et al., 2003). GR and APX activity was determined according to the methods described by Lin et al. (2018) and Kuo et al. (2020b), respectively. Protein concentrations were quantified using the Coomassie Blue dye binding method (Bradford, 1976) using a concentrated dye purchased from BioRad (500-0006, Hercules, CA, USA). AsA and DHA concentrations were determined by an ascorbate oxidase based method according to Lin et al. (2016). GSH and oxidized glutathione (GSSG) were extracted from the frozen algal cell pellet (obtained from 5 mL samples) using 5% (w/v) trichloroacetic acid (TCA) and determined at 412 nm according to Lin et al. (2018).

### Determination of NADPH oxidase activity

NADPH oxidase activity was determined according to Kaundal et al.’s (2012) method based on O_2_^−.^ production. After homogenization, centrifugation at 10,000 ×*g*, and then ultra-centrifugation at 203,000 ×*g* for collection of total membranes, the pellet was suspended in 1 mL ice-cold 10 mM Tris-HCl (pH 7.4). The reaction mixture consisted of 50 mM Tris-HCl buffer (pH 7.5), 1 mM XTT (3-[1-[phenylamino-carbonyl]-3,4-tetrazolium]-bis(4-methoxy-6-nitro)benzenesulfoni c acid hydrate), 1 mM NADPH, and 5 μg membrane protein. The reduction rate of XTT by O_2_^−.^ detected at 492 nm was used to calculate enzyme activity; the extinction coefficient used was 2.16 × 10^4^ M^−1^ cm^−1^.

### RNA Isolation, cDNA Synthesis, Transcriptomic Analysis, and mRNA Quantification via Real-Time Quantitative PCR

Total RNA was extracted using the TriPure Isolation Reagent (Roche Applied Science, Mannheim, Germany) according to the manufacturer’s instructions. The methods for cDNA library preparation, Illumina sequencing, and sequence analysis are described in Supplemental Fig. S6. For qPCR assay, the total RNA concentration was adjusted to 2.95 μg total RNA·μL^−1^ and treated with DNase (TURBO DNA-free^TM^ Kit, Ambion Inc., The RNA Company, USA) to remove residual DNA. Then 1.5 μg of total RNA was used for the preparation of cDNA. cDNA was amplified from the poly-(A+) tail using Oligo (dT)12-18 with the VersoTM cDNA Kit (Thermo Fisher Scientific Inc., Waltham, MA, USA), and the volume was adjusted to a concentration of 30 ng·mL^−1^ based on original RNA quantity in each sample. The primers for the targeted genes are listed in Supplemental Table S11. The real-time quantitative PCR was performed using a LightCycler 480 system (Roche Applied Science, Mannheim, Germany). A PCR master mix was prepared with the LightCycler 480 SYBR Green I Master Kit (Roche Applied Science, Mannheim, Germany). Each reaction was performed in a total volume of 10 μL, containing 1× LightCycler 480 SYBR Green I Master Mix, the selected concentration of each primer, and cDNA corresponding to 30 or 50 ng·μL^−1^ RNA in the reverse transcriptase reaction. The amplification program consisted of an initial denaturation at 95°C for 5 min, followed by 50 amplification cycles of annealing at 60°C for 10 s, elongation at 72°C for 5 s, real-time fluorescence measurements, and finally, denaturation at 95°C for 15 s. The 2-^ΔΔ^CT method was used to calculate the relative change in mRNA level normalized to a reference gene (UBC, NCBI: AY062935) and the fold increase was calculated relative to the control RNA sample at 0 min. Because the results based on the EF-1α internal control were similar to those based on UBC, the relative changes in the levels of gene transcription were expressed based on UBC.

### Western Blots

Soluble protein was extracted according to Kuo et al. (2020a). For each sample, 30 μg of protein was loaded into each lane, resolved on a 15% SDS-PAGE gel, and transferred to a polyvinylidene fluoride membrane for antibody binding with rabbit polyclonal anti-ATG8 antibody (ab77003; Abcam, Cambridge, UK), anti-CrDHAR (LKT BioLaboratories Ltd., Taoyuan, Taiwan), or a mouse monoclonal antibody against α-tubulin (ab11304; Abcam, Cambridge, UK). Following the incubation with horseradish peroxidase-conjugated secondary antibodies (MD20878; KPL, Gaithersburg, MD, USA), the immunoblots were visualized and the relative abundance of ATG8, ATG8-PE, or CrDHAR1 protein was estimated based on α-tubulin intensity.

### Statistics

Three independent biological replicates were performed and all experiments were repeated at least three times. Because the replications showed similar trends, only the results from one replicate were shown in this paper. Statistical analyses were performed using SPSS (SPSS 15.0, Chicago, IL, USA). Significant differences between means were analyzed using Student’s *t*-test or Scheffe’s test following significant analysis of variance for the controls and treatments at *P* < 0.05.

### Accession Numbers

The transcriptome sequences can be accessed from the Sequence Read Archive (SRA) website using the BioProject accession number: PRJNA629395 (https://www.ncbi.nlm.nih.gov/books/NBK158898/) and the BioSample accessions SAMN14775313, SAMN14775314, SAMN14775315, SAMN14775316, SAMN14775317, SAMN14775318, SAMN14775319, SAMN14775320, SAMN14775321, and SAMN14775322.

## Supplemental Data

The following supplemental materials are available:

Supplemental Tables S1–S11.

Supplemental Figures S1–S10.

## ACKNOWLEDGEMENTS

We thank Dr. Su-Chiang Fang, Academia Sinica Biotechnology Center in Southern Taiwan, for assistance with culturing *Chlamydomonas* and manuscript preparation.

